# Glandular architecture and malignant behaviour in colorectal cancer is regulated by the sialomucin Podocalyxin

**DOI:** 10.64898/2026.07.10.737619

**Authors:** Erin M. Cumming, Kai Rakovic, Kathryn AF. Pennel, Laura A. Galbraith, Emma Sandilands, Louise Mitchell, Lynn McGarry, Rene Jackstadt, Kathryn Gilroy, Colin Nixon, Owen J. Sansom, John Le Quesne, Karen Blyth, Joanne Edwards, David M. Bryant

## Abstract

Glandular architecture-the coordination of lumen-containing structures by an apical-basal polarised epithelium-is frequently maintained in colorectal cancer (CRC), yet whether it actively contributes to tumour progression or metastatic competence remains unclear. Here, we identify Podocalyxin (PODXL), a developmental regulator of epithelial lumen formation, as a key determinant of glandular tumour architecture in CRC. PODXL is upregulated in CRC, particularly in poor-prognosis Consensus Molecular Subtype 4 (CMS4) tumours, where high expression predicts reduced survival. Using genetically engineered mouse models, matched organoids, human cell lines and xenografts, we show that PODXL promotes organisation of CRC cells into gland-like, lumen-containing structures. Loss of PODXL disrupts glandular architecture in both primary tumours and liver metastases, reducing tumour growth and metastatic colonisation. Mechanistically, TGF-β signalling drives PODXL upregulation. Together, these findings establish glandular architecture as an active determinant of CRC progression and identify PODXL as a functional contributor rather than merely a prognostic biomarker.

## Introduction

Glandular architecture is a defining histopathological feature of colorectal cancer (CRC). Whether this epithelial organisation - including apical-basal polarity and lumen formation - actively contributes to tumour progression or instead represents a vestigial feature progressively lost during malignant evolution remains unclear. Epithelial polarity and tissue organisation are traditionally considered tumour-suppressive, with their disruption promoting tumourigenesis^1,2^. However, many epithelial tumours retain organised gland-like structures, and emerging evidence suggests that epithelial architecture can be maintained and co-opted to support tumour growth^3,4^. This raises a fundamental question: whether glandular architecture merely reflects tumour differentiation, or whether it actively contributes to malignant progression.

Tumours frequently co-opt developmental and stem-like programmes during progression^5–9^, raising the possibility that the developmental mechanisms governing epithelial organisation are similarly redeployed during tumourigenesis.

Podocalyxin-like 1 (PODXL) is a transmembrane sialomucin that plays a central role in epithelial polarity and, consequently, lumen formation during tissue morphogenesis. It belongs to the CD34-family, characterised by a dense array of negatively charged glycans in the extracellular domain^10–12^. This extensive glycosylation generates electrostatic repulsion between adjacent membranes^12,13^, facilitating physical separation of neighbouring cells and establishing where the apical lumen forms^14–18^. In contrast to its extracellular domain, the PODXL intracellular domain is highly conserved across species. The intracellular domain contains binding sites for scaffold proteins^12,19^, most notably NHERF1/2^20,21^ and Ezrin, which link PODXL to the actin cytoskeleton^21,22^ and enable it to function as a signalling platform for Rho-family GTPase-dependent cytoskeletal remodelling^21,22^. Through these interactions, PODXL integrates membrane identity with cytoskeletal organisation, coupling apical membrane specification to downstream pathways that regulate epithelial cell shape, polarity, and lumen assembly in development. Notably, although PODXL is essential for proper podocyte morphogenesis in the kidney – where genetic deletion results in perinatal lethality – it is not universally required for apical-basal polarity across all epithelial tissues^23,24^. PODXL is expressed in multiple organs, including vascular endothelium, haematopoietic progenitors, mesothelium and subsets of neurons^12,23,25^, yet its loss outside the kidney does not produce comparable global polarity defects. Epithelial polarity networks are highly modular and buffered by multiple interacting protein complexes^2,26,27^, consistent with tissue-specific compensatory mechanisms operating in many organs. In contrast, tumour cells that retain or co-opt epithelial architecture may acquire a selective dependence on PODXL-mediated structural organisation.

Consistent with this, PODXL expression is frequently elevated across multiple cancer types, where it is associated with aggressive disease and poor patient survival^28–32^. Despite its established developmental role in epithelial polarity, PODXL expression in some tumour cells correlates with mesenchymal markers such as Vimentin and N-cadherin and is upregulated during TGF-β–induced epithelial–mesenchymal transition (EMT)^33,34^. Part of this mechanism, at least in prostate cancer cells, involves attenuated PODXL ubiquitination - which normally promotes PODXL removal from the cell surface - in mesenchymal-state cells, thereby increasing cell-surface-associated functions of PODXL^35^. Functionally, PODXL has been reported to enhance cell motility, invasion and metastatic dissemination through Ezrin-dependent regulation of actin dynamics and Rho GTPase activity^34^, as well as by acting as a decoy receptor for the invasion-suppressing Galectin-3 protein to promote integrin-dependent invasion^35^. In CRC, PODXL expression is commonly elevated and independently associated with poor prognosis^28–30,36^. However, existing studies have largely identified PODXL as a biomarker of aggressive disease without establishing whether it functionally promotes CRC growth, architectural organisation or metastatic competence.

CRC provides an ideal system in which to address this question because tumour evolution occurs within a gland-forming epithelium where tissue organisation remains a defining histopathological and clinical feature. Although tumour progression is often associated with loss of epithelial polarity and differentiation - and tumour grading routinely incorporates assessment of glandular architecture - it is increasingly recognised that epithelial organisation is not uniformly tumour suppressive, and that specific organisational states can actively promote malignancy^37–40^. Notably, McNagny, Roskelley and colleagues demonstrated that PODXL overexpression in breast cancer cells induces organised, lumen-containing structures that facilitate cohesive invasion^41^, raising the possibility that tumour cells co-opt apical-basal polarity programmes to promote progression. Together, these observations prompted us to ask whether PODXL’s conserved developmental role in epithelial lumen formation is redeployed within CRC to organise tumour architecture, and whether this organisation contributes functionally to malignant progression.

Here, we examine PODXL expression and function across CRC patient cohorts, genetically engineered mouse models, matched organoids, human cell lines and xenografts. We identify TGF-β signalling as a key upstream regulator of PODXL elevation and demonstrate that PODXL promotes organisation of tumour cells into gland-like, lumen-containing structures both in vitro and in tumours. Loss of PODXL disrupts glandular architecture across primary and metastatic sites and reduces metastatic burden. Together, these findings demonstrate that glandular architecture is an active determinant of CRC progression and identify PODXL as a developmental epithelial morphogenesis regulator co-opted to support tumour growth and metastatic colonisation.

## Results

### PODXL expression is elevated in colorectal cancer and predicts poor survival in CMS4 tumours

Elevated PODXL protein has been associated with poor prognosis in CRC, based almost exclusively on immunohistochemical (IHC) analyses in selected patient cohorts^28–30,36^. However, the association between *PODXL* mRNA levels and disease characteristics or prognostic outcome has not previously been systematically examined. To determine whether elevated *PODXL* expression at the transcript level represents a consistent and clinically relevant feature of CRC, we analysed *PODXL* mRNA expression across multiple independent patient cohorts.

In The Cancer Genome Atlas (TCGA) dataset^42^, *PODXL* mRNA was significantly higher in the tumour compared to normal tissue, both when colon (colon adenocarcinoma, COAD) and rectal (rectal adenocarcinoma, READ) cancers were analysed together (Fig. 1A) and when examined separately (Fig. S1A–B). This increase was also observed when the analysis was restricted to matched normal-tumour pairs from individual patients (Fig. 1B, S1C-D). This association was independently replicated across 10 additional cohorts profiled using alternative array platforms (Fig. S1E-N). *PODXL* copy-number alterations did not correlate with mRNA levels (Fig. S1O), suggesting that elevated *PODXL* mRNA expression is driven primarily by transcriptional regulation rather than genomic amplification.

**Figure 1.**
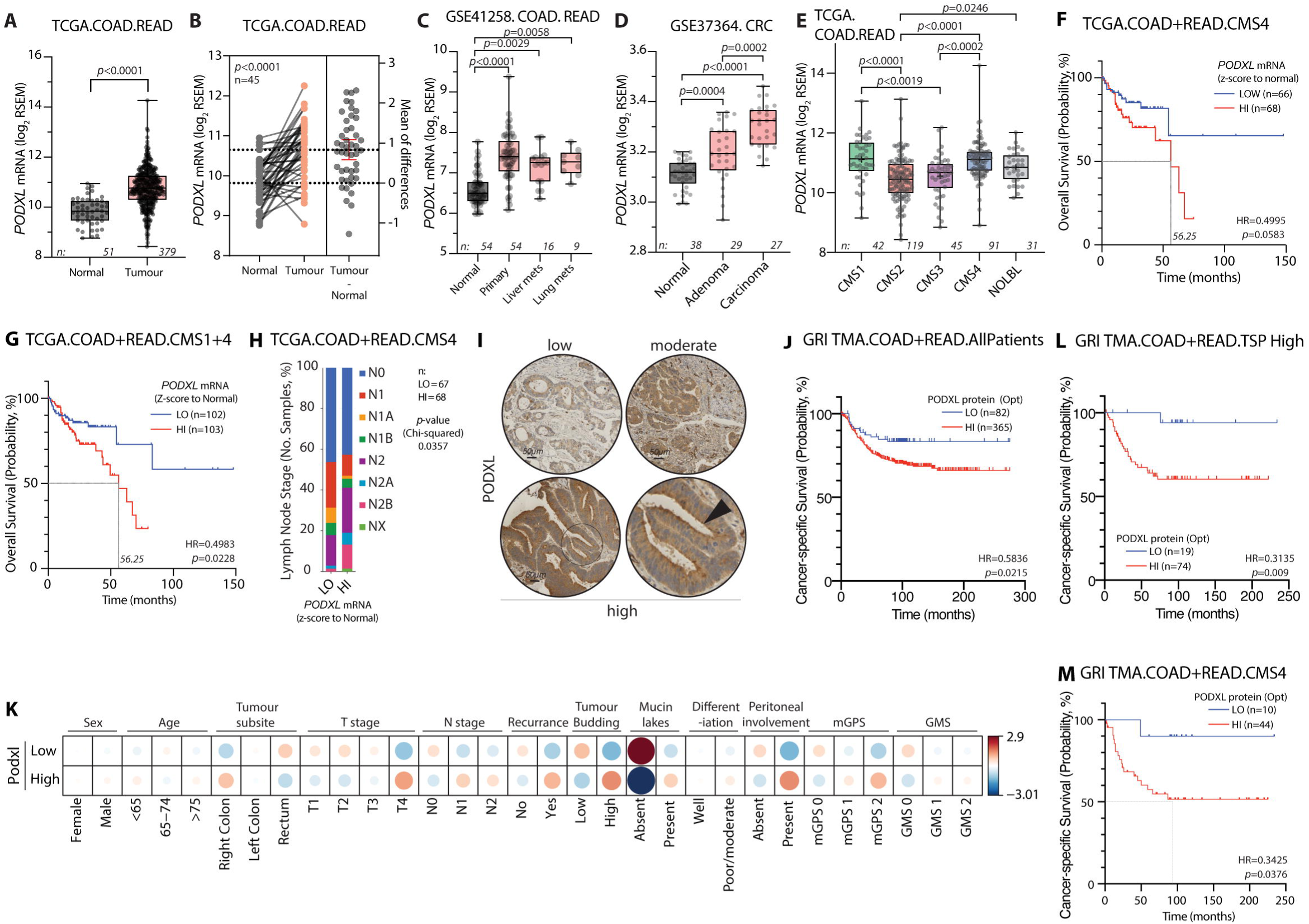
*PODXL* expression is elevated in CRC and predicts poor survival in CMS4 patients. **A-E,** *PODXL* mRNA expression in: **(A)** unmatched normal versus tumour tissue, TCGA (PanCancer Atlas 2018) combined colon and rectal adenocarcinoma (COAD + READ), **(B)** matched normal–tumour pairs from the same patients, TCGA COAD + READ, **(C)** normal tissue, primary tumours and metastases, (COAD + READ; GEO), **(D)** normal tissue, adenoma and carcinoma, (CRC), **(E)** across Consensus Molecular Subtypes (CMS1–4), TCGA COAD + READ. NOLBL, no label (unclassifiable). **F,G.** Overall survival stratified by median-based *PODXL* mRNA in **(F)** CMS4 or CMS1+4 **(G)** patients, TCGA COAD + READ. **H,** Lymph-node (N) stage distribution in CMS4 patients stratified by *PODXL* expression (TCGA COAD + READ, stacked bar). **I,** Representative PODXL immunohistochemistry (low, moderate, high) of the CRC TMA; PODXL labelling localises to the apical/luminal domain of epithelia in PODXL-high tissue (arrowhead and circle). **J,** Cancer-specific survival stratified by PODXL protein level (optimised cutoff) in the TMA cohort for **(K)** all patients. **K,** Associations between PODXL protein expression (Low vs High, optimized cut off) and clinicopathological features in the TMA cohort: sex, age, tumour subsite, T stage, N stage, recurrence, tumour budding, mucin lakes, differentiation, peritoneal involvement, mGPS (modified Glasgow Prognostic Score) and GMS (Glasgow Microenvironment Score) (also see Table S1). Circle SIZE = absolute value of the Chi-squared residuals and COLOUR = sign of the residual (red positive, blue negative); the numerical state at the edge = Chi-squared residual values (which is why it can exceed - /+1). **L,M,** Cancer-specific survival stratified by PODXL protein (optimised cutoff) in the TMA cohort for **(L)** patients with high tumour–stroma percentage (TSP), or **(M)** CMS4 TMA patients. *Box-and-whisker plots (A, C, D, E): midline, median; box, IQR; whiskers, range; dots, individual datapoints; Mann–Whitney two-tailed test. Estimation plot (B): paired t-test. Kaplan–Meier survival (F, G, J, L, M): log-rank (Mantel–Cox) test, hazard ratios shown. H: chi-squared test. Datasets/accession numbers, patient sample numbers and tests are indicated in each panel*.

This robust, reproducible tumour-specific increase across independent cohorts prompted us to ask whether *PODXL* upregulation tracks with disease progression. Comparison of primary tumours with liver or lung metastases across two independent datasets revealed that while *PODXL* mRNA was elevated in primary tumours relative to normal tissue (Fig. S1P, Q), it was not consistently further increased in metastatic lesions from these same datasets (Fig. 1C, S1R-T). Consistent with this, *PODXL* mRNA expression did not increase with tumour, node, metastasis (TNM) stage across four independent cohorts (Fig. S1U-X). This indicates that *PODXL* upregulation is an early, stage-independent feature of colorectal tumours rather than a marker of advanced disease.

To investigate earlier events in colorectal tumourigenesis, we analysed *PODXL* expression in datasets including pre-malignant adenomas. *PODXL* mRNA expression increased stepwise from normal tissue to adenoma to carcinoma (Fig. 1D). Similarly, *PODXL* mRNA expression was lowest in normal epithelial compartments (crypt and surface epithelium), then increased in adenomas and carcinomas (Fig. S1Y). In paired analyses, *PODXL* mRNA expression was higher in adenomas compared to matched normal tissue (Fig. S1Z). Although the adenoma-to-carcinoma increase was significant in one cohort (Fig. 1D) but not the other (Fig. S1Y), the overall pattern was consistent with progressive *PODXL* upregulation during early tumourigenesis. Together, these data suggest that *PODXL* mRNA upregulation may be initiated early during colorectal tumourigenesis, emerging at the adenoma stage and maintained through carcinoma and metastasis.

We next examined whether *PODXL* mRNA expression may differ between different CRC molecular subtypes. CRCs can be classified into four Consensus Molecular Subtypes (CMS1–4) that differ in their molecular features, biological behaviour, and clinical outcomes^43^. Among these, CMS4 tumours - characterised by stromal infiltration, mesenchymal features, and elevated TGF-β pathway activity - are associated with the poorest prognosis^43^. Across seven independent patient cohorts, *PODXL* mRNA expression was consistently highest in CMS1 and CMS4 tumours relative to CMS2 and CMS3 (Fig. 1E, Fig. S1AA-AF). The elevation in CMS4 is notable given its association with the worst clinical outcomes, while high expression in CMS1 - the microsatellite-unstable, immune-active subtype - may in part reflect *PODXL*’s known expression in immune and endothelial cell populations within the tumour microenvironment.

We examined whether *PODXL* mRNA expression levels were associated with patient survival, and whether this relationship differed according to molecular subtype. Across all patients, a median split of *PODXL* mRNA expression did not significantly stratify overall survival (Fig. S2A). However, when patients were stratified by CMS subtype, *PODXL* mRNA significantly stratified survival of patients with CMS4 profile tumours showing lower overall survival (Fig. 1F). Combining CMS1 and CMS4 patients further strengthened this association (Fig. 1G). Elevated *PODXL* in CMS4 patients was associated with higher lymph node staging, consistent with increased metastatic spread (Fig. 1H). In contrast, CMS2 survival curves for high and low *PODXL* crossed over time (Fig. S2B), indicating a non-proportional association without a consistent directional effect. CMS3 and CMS1 patients showed no significant stratification by *PODXL* levels alone (Fig. S2C, D). This indicates that the elevation of *PODXL* in CMS4 patients is associated with poor outcome.

To validate these transcript-level findings at the protein level, we performed PODXL IHC on a tissue microarray (TMA) comprising CRC tissue from 447 stage II–III patients. A median split of all patients based on PODXL labelling, similar to *PODXL* mRNA, did not stratify survival (p=0.0943, Fig. S2G). However, using an optimised scoring approach and a data-driven cut-point (Methods), patients were stratified into PODXL-low and PODXL-high groups (Fig. 1I). Here, high PODXL protein expression associated with significantly decreased survival (Fig. 1J). In high-expression tissues, PODXL labelling was localised to the apical luminal domain of epithelia (Fig. 1I). High PODXL protein expression was not significantly associated with baseline clinicopathological characteristics (Fig. 1K, Table S1), including sex (p=0.935), age (p=0.886), tumour subsite (p=0.288), or differentiation status (p=0.722). Nodal status (p=0.213) and recurrence (p=0.128) were also not significantly associated with PODXL expression (Fig. 1K, Table S1). However, high PODXL expression was significantly associated with features of aggressive tumour behaviour (Fig. 1K), including increased tumour budding (high PODXL, 35.9% vs low PODXL, 25.7%, p=0.021), mucin lakes (40.6% vs 19.5%, p=0.002), and peritoneal involvement (28.1% vs 19.6%, p=0.034). A trend towards right-sided presentation and more advanced T stage (T4) was noted in the high PODXL protein group (Fig. 1K, Table S1), which was corroborated in TCGA data for *PODXL* mRNA expression (Fig. S2E). Consistent with this, high PODXL mRNA was associated with more advanced local tumour (T) stage in TCGA (Fig. S2F); this reflects an association with local invasion depth (T) rather than with overall TNM stage, which did not track with PODXL (Fig. S1U–X). No significant associations were observed with systemic inflammatory scores (Fig. 1K, Table S1), including the modified Glasgow Prognostic Score (mGPS, p=0.112) - a measure of the systemic inflammatory response based on serum C-reactive protein (CRP) and albumin levels - or the Glasgow Microenvironment Score (GMS, p=0.441)^44^, derived from tumour budding and peritumoural inflammatory infiltrate assessed using the Klintrup–Mäkinen grade^45^.

Supporting a role for PODXL in aggressive tumour phenotypes, PODXL expression significantly stratified survival in patients with a low, but not high, Klintrup–Mäkinen score - a measure of local inflammatory cell infiltrate at the invasive margin of the tumour (Fig. S2H,I) - suggesting that the effects of PODXL may be most evident in immunosuppressed tumours. Additionally, PODXL protein was significantly associated with reduced survival in tumours with a high, but not low, tumour-stroma percentage (TSP) (Fig. 1L, Fig. S2J), indicating that PODXL-associated effects occur in stroma-rich tumours. Consistent with this, and mirroring findings for *PODXL* mRNA expression, stroma-rich, immune-infiltrate-low CMS4 tumours showed the strongest association between high PODXL protein expression and poor survival (Fig. 1F,M), whereas other molecular subtypes showed no significant stratification (Fig. S2K–M). Unlike *PODXL* mRNA expression, combining CMS1 and CMS4 tumours reduced the significance of survival stratification for PODXL protein labelling (Fig. S2N). Together, these data establish *PODXL*, at both transcript and protein level, as a subtype-specific prognostic marker in CRC, with its strongest predictive value in stroma-rich, immune-suppressed CMS4 tumours - prompting us to turn to experimental models to ask how *PODXL* is regulated and whether it actively shapes tumour architecture.

### PODXL localises to the apical domain of glandular tumour epithelium in CMS4 mouse tumours

Given the enrichment of PODXL expression in poor-prognosis human CRC patient CMS4 tumours, which are characterised by prominent stromal activation and elevated TGF-β signalling^43^, together with prior reports demonstrating TGF-β–mediated induction of PODXL expression in lung cancer cells^33^, we hypothesised that TGF-β signalling may contribute to PODXL upregulation in CRC. To investigate this, we first examined *Podxl* expression in genetically engineered CRC mouse models (GEMMs) that recapitulate CMS4-like features (Fig. 2A)^46^. The first CMS4 model, designated ‘KP’, combines oncogenic *Kras* (*Kras^G12D/+^*; ‘K’) and loss of the tumour suppressor *Trp53* (*Trp53^-/-^*; ‘P’) alleles. The second, designated ‘KPN’ additionally incorporates constitutive activation of Notch signalling (*Notch1-Intracellular domain*; *Notch1^ICD-IRES-nEGFP^*; ‘N’) to generate aggressive, metastatic colorectal tumours (Fig. 2A)^46^. Both models have been independently classified as CMS4-like based on transcriptomic profiling using the MmCMS classifier^46,47^. While both models exhibit CMS4-like molecular features, KP tumours rarely metastasise, whereas KPN tumours – which possess Notch-driven elevated TGF-β pathway activity - frequently spread to the liver and extrahepatic sites^46^. The A+K genotype is included for comparison as a genotype not able to be classified by CMS.

**Figure 2.**
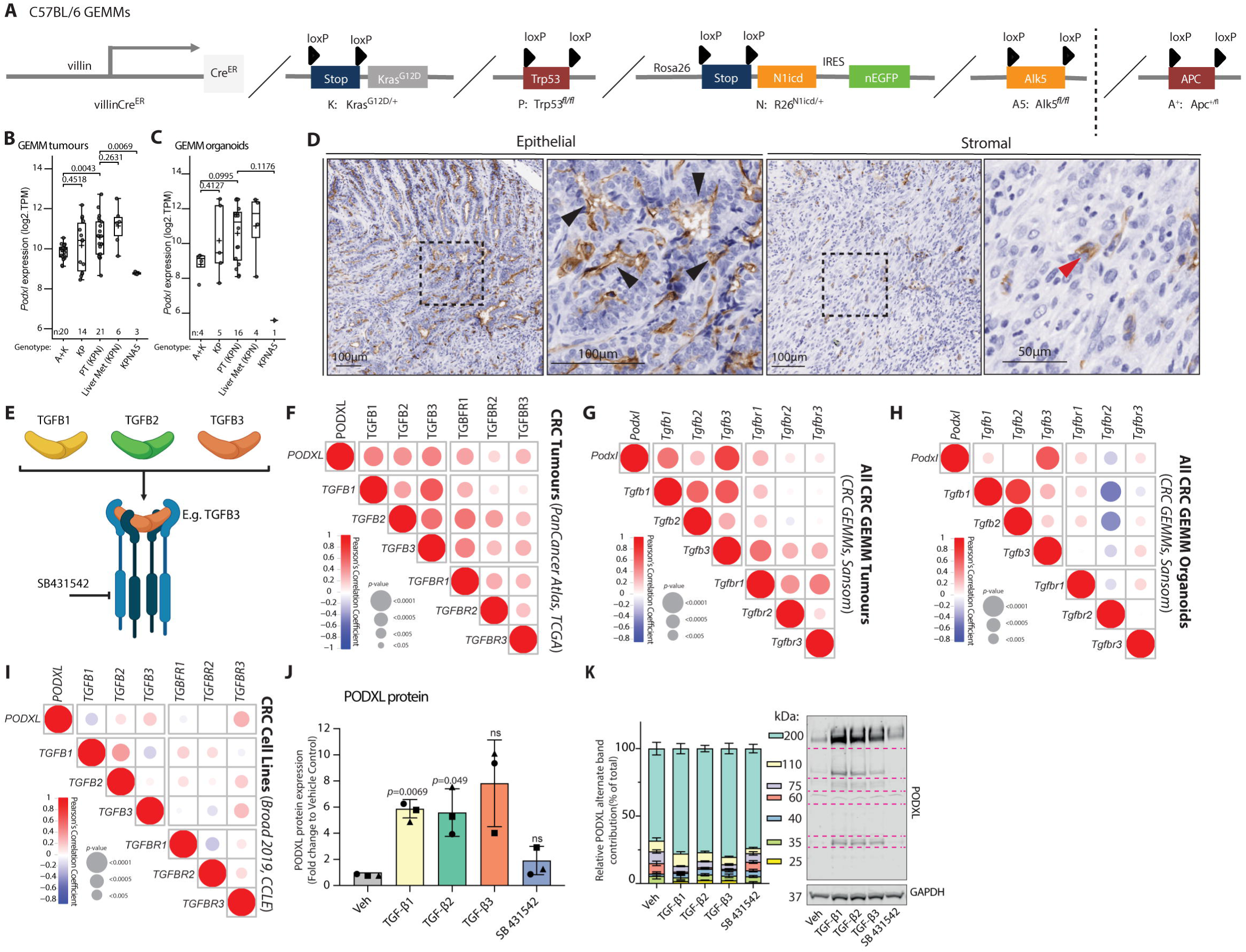
TGF-β signalling is associated with elevated PODXL expression in CMS4 colorectal cancer. **A,** Schematic of the villin-CreER–driven Cre/loxP conditional alleles used to generate the CMS4-like CRC GEMMs: Kras^G12D/+ (K), Trp53^fl/fl (P), Rosa26^N1icd/+ (N) and Alk5^fl/fl (A5). The dashed vertical line separates these from Apc^+/fl (A+), which was combined with Kras^G12D/+ (K) to generate the A+K genotype - an adenoma-forming, non-CMS definable comparator. **B, C,** *PODXL* mRNA levels assessed by RNA-seq in GEMMs **(B)** primary tumours (PT) and liver metastases or **(C**) in GEMM-derived organoids from indicated genotypes and tissues. Data are from the Gold Standard set, 2026 Sansom lab. Sample numbers for organoids indicate individual organoid lines. **D,** Immunohistochemical staining of PODXL in KPN tumours showing predominant apical/luminal localisation in tumour glandular epithelium (black arrowheads) and a minor pool of stromal staining (red arrowheads). **E,** Schematic of canonical TGF-β signalling. TGFB1, TGFB2 and TGFB3 ligands engage TGFBR1/TGFBR2 heterodimers, with TGFB3 as the example ligand, to activate downstream signalling. SB-431542, a small-molecule inhibitor of TGFBR1 kinase activity. **F–I,** Pearson correlation matrices showing associations between *PODXL/Podxl* mRNA and TGF-β ligand and receptor mRNA across CRC datasets: human TCGA PanCancer Atlas CRC tumours (F), CRC GEMM primary tumours (G; Sansom lab, 2020), CRC GEMM-derived organoids (H; Sansom lab, 2020), and human CRC cell lines (I; CCLE, Broad 2019). Pearson correlation coefficients (r) *with two-tailed t-test* are shown on a blue-to-red scale; dot size indicates p-value. **J,** PODXL protein expression in HT-29 cells 24 h after treatment with TGF-β1, TGF-β2, TGF-β3 or the TGFBR1 inhibitor SB-431542, quantified by western blot as PODXL normalised to GAPDH loading control and presented as fold change relative to vehicle control. Data shown from three independent experiments. **K,** Representative western blot of PODXL in HT-29 cells from one of three experiments across treatment conditions, showing multiple molecular-weight species of which the ∼200 kDa band corresponds to the mature, high-molecular-weight PODXL glycoform; left panel, stacked-bar quantification of each band’s contribution to the total PODXL signal. GAPDH, loading control. *Box-and-whisker plots (B, C): midline, median; box, IQR; whiskers, range; Bar plot (J): box, mean, ± SD dots, individual datapoints; Stacked plot (K) boxes mean* ± SEM.

Quantitative analysis of bulk RNA-seq demonstrated that *Podxl* mRNA was significantly elevated in the Notch/TGFB-driven CMS4-profile KPN primary tumours compared with the non-CMS4 A+K comparator (Fig. 2B; p=0.0043), whereas non-Notch/TGFB-driven KP tumours showed intermediate expression that was not significantly distinct from A+K (Fig. 2B; p=0.45). This elevation was blunted in KPN mouse derived organoids ex vivo, indicating a microenvironment contribution to *Podxl* expression; in organoids neither genotype differed significantly from the A+K comparator, though trended in the same direction (Fig. 2C; KPN vs A+K, p=0.0995; KP vs A+K, p=0.4127). Consistent with the patient expression data (Fig. 1D), *Podxl* mRNA was not further elevated in KPN liver metastases relative to primary tumours, nor in KPN organoids derived from these sites, reinforcing a pattern of upregulation that is maintained through metastatic progression. IHC analysis of KPN tumours showed that, as in CRC patient tissue (Fig. 1I), PODXL protein localised predominantly to the apical surface of tumour epithelial cells lining glandular structures (Fig. 2D, black arrowhead), consistent with its established role in epithelial polarity and lumen morphogenesis^16,17^, with a minor pool of PODXL-positive cells observed in the stroma of some regions (Fig. 2D, red arrowheads). Together, these data in GEMM primary tumours, and to a lower extent GEMM-derived organoids, mirror the observations in CRC patients, with *Podxl* mRNA and PODXL protein both upregulated in CMS4 CRC.

### TGF-β elevates PODXL expression

As Podxl elevation in KPN tumours was attenuated in matched organoids ex vivo, we reasoned that a microenvironmental signal - most likely TGF-β, the defining feature of the CMS4 subtype - drives its upregulation, and set out to test this directly. Canonical TGF-β signal transduction occurs via coupling of any of TGF-β1-3 ligands with obligate signalling through TGFBR1/2 heterodimers (Fig. 2E), which can be inhibited by the small molecule SB-431542^48^. In line with TGF-β signalling being a major driver of the CMS4 phenotype, primary tumours from KPN models with genetic deletion of the TGFBR1 (*Alk5*, KPNA5) exhibited significantly reduced, but not abolished, *Podxl* mRNA expression (Fig. 2B). A single KPN organoid line lacking TGFBR1 (KPNA5) also showed reduced *Podxl* mRNA, although this observation is limited to one sample (Fig. 2C).

To extend these findings beyond the CMS4 GEMM system and assess the generality of the TGF-β–*PODXL* relationship, we performed correlation analyses across multiple independent human CRC mRNA datasets. *TGFB3* mRNA expression exhibited a robust and consistent correlation with *PODXL* mRNA expression across CRC patient tumours ^42^, GEMMs, and GEMM-derived organoids (Fig. 2F-H). Notably, this correlation was observed without sub-setting to CMS subtypes and is therefore not restricted to CMS4 tumours. In contrast to the ligand *TGFB3*, the TGF-β receptors (*TGFBR1–3*) showed no consistent correlation with *PODXL* across datasets; *TGFBR1* correlated modestly in patient tumours (Fig. 2F) but not in the GEMM or cell-line datasets (Fig. 2G–I). These analyses identify a conserved link between *PODXL* and TGF-β ligand signalling across patient tumours, GEMMs and organoids; we next asked whether this relationship was preserved in cultured CRC cell lines.

In cultured cell lines this association was more modest (Fig. 2I), mirroring the attenuation seen *ex vivo* in GEMM-derived organoids relative to tumours. Across a panel of 57 human CRC cell lines, we assessed in silico whether *PODXL* mRNA expression was related to origin (primary versus metastasis), dependency in CRISPR screens from DepMap as an indicator of growth regulation, or any of the most frequent human CRC mutations (Fig. S3A). In contrast to patient tumours, GEMMs and GEMM-derived organoids, no association between *PODXL* mRNA levels and CMS classification was observed (Fig. S3A, B), and this was unrelated to whether cell lines were derived from primary tumour or metastases (Fig. S3A, C). Using DepMap dependency scores to indicate potential growth defects upon *PODXL* knockout (KO), no major differential dependency on *PODXL* loss was observed based on *PODXL* mRNA levels (Fig. S3A, D), by CMS subdivision (Fig. S3A, E; though note significant but minor difference between CMS2 and CMS4) or whether cell line was derived from primary tumour or metastasis (Fig. S3A,F).

In probing a selection of CRC cell lines from different CMS, PODXL protein was detected as multiple molecular-weight species (glycoforms) (Fig. S3G-I), did not correlate with *PODXL* mRNA abundance (Fig. S3J). Together, these data indicate that the robust association between PODXL expression and CMS4 observed in vivo is blunted in ex vivo cultured organoids (Fig. 2C) and largely lost in long-term cultured CRC cell lines (Fig. S3). This suggests that additional extrinsic cues, such as CMS4-associated TGF-β signalling, may be required to sustain elevated PODXL expression in vivo. We therefore examined whether exogenous TGF-β addition could elevate PODXL expression, with TGF-β3 predicted as the potential key inducer. We treated TGF-β-competent HT-29 cells individually with each ligand for 24 hours, to determine any potential differences in ligand-induced signalling. TGF-β1 and TGF-β2 significantly increased, and TGF-β3 also increased (though not significantly), PODXL protein (Fig. 2J), most prominently affecting the mature high-molecular-weight (∼200 kDa) PODXL glycoform (Fig. 2K). Given that TGFBR1 (Alk5) knockout in murine CRC CMS4 tumours and organoids decreased, but did not eliminate, *Podxl* mRNA expression (Fig. 2B, C), we asked whether TGF-β signalling is required for baseline PODXL expression or only results in its elevation. Neither chemical inhibition of TGFβR1 with SB-431542^48^ (Fig. 2J-K), or shRNA-mediated TGFβR1 depletion (Fig. S3K-M) in HT-29 cells reduced baseline PODXL. Together, these findings indicate that TGF-β signalling elevates PODXL expression above baseline levels in CRC human cells and murine tumours and organoids, but that TGF-β signalling is dispensable for basal PODXL expression. Moreover, this association is attenuated in cells cultured ex vivo but can be restored by exogenous TGF-β ligand addition.

### PODXL controls collective organisation and lumen formation in 3D culture

We next examined the functional consequence of PODXL loss in CRC cells. We generated two independent PODXL knockout (KO) HT-29 cell lines (KO1, KO2) alongside a scrambled-sequence control (SCR) using CRISPR-Cas9 (Fig. 3A). As the sgPODXL-2 line (KO2) gave the greatest reduction in PODXL expression, it was used for all subsequent experiments unless KO1 is also indicated. Mirroring the lack of a robust growth dependency on PODXL in large-scale CRISPR screens of CRC cells in vitro (Fig. S3E,F), PODXL KO did not alter growth in either colony-formation or two-dimensional growth assays (Fig. S4A–D).

**Figure 3.**
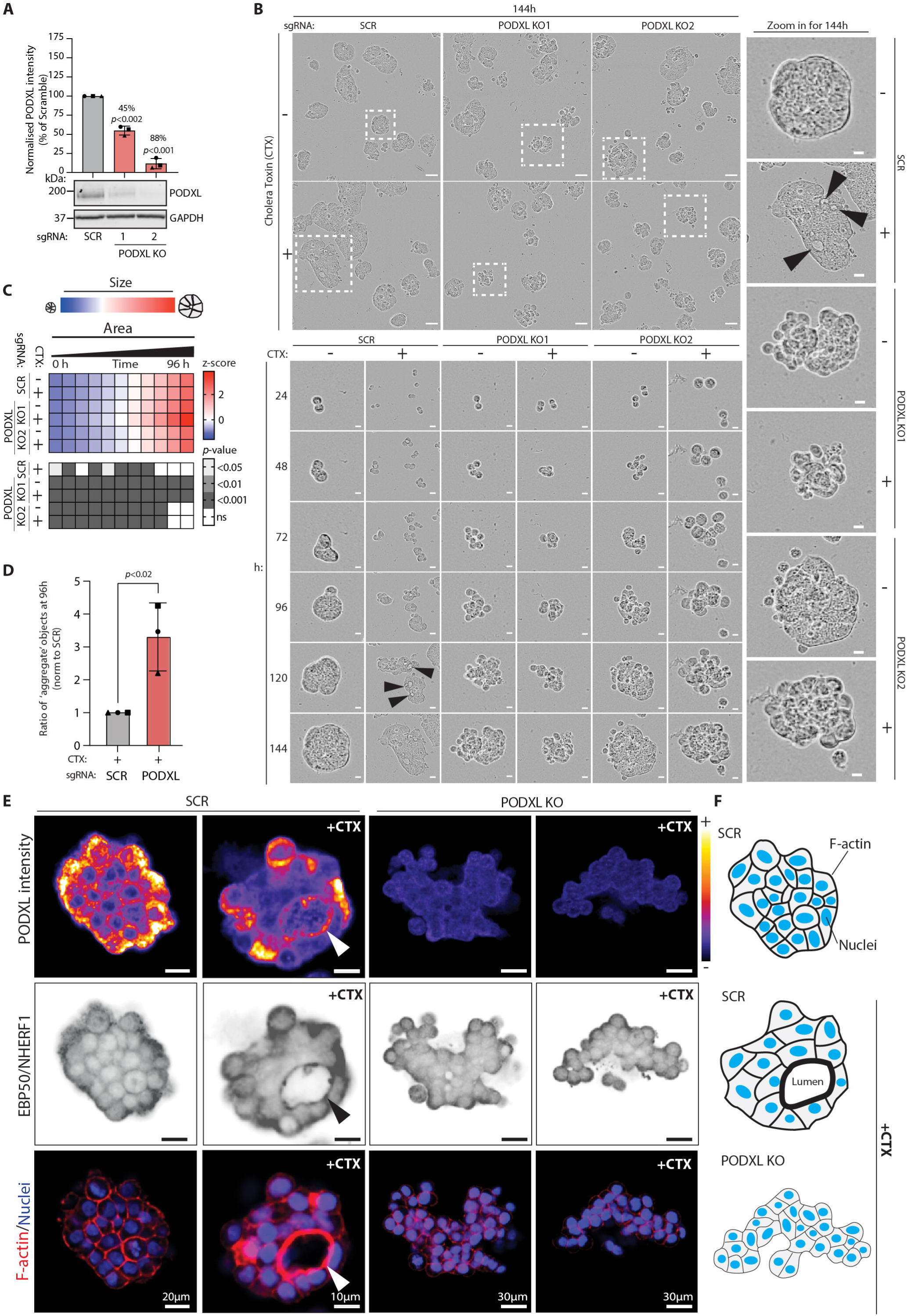
PODXL controls collective organisation and lumen formation in 3-dimensional culture. **A,** Immunoblot confirming PODXL knockout in HT-29 cells (sgRNA scramble or two independent *PODXL* sgRNAs: KO1, KO2), GAPDH loading control with quantification. Datapoint shapes correspond to each of 3 independent experimental replicates. **B,** Representative phase-contrast images from live imaging of control (SCR) and PODXL KO HT-29 spheroids grown with or without cholera toxin from 3 independent experiments, each containing 4 technical replicates. Arrowheads, lumens. Note that images on right labelled ‘zoom in for 144h’ are the same images as in bottom row (144h) but presented in larger format for ease of visualising lumens. Scale bars 95 µm. **C-D,** Analyses if all experimental images corresponding to Fig. 3B, not only images shown. **(C)** Heatmap quantification of spheroid (as indicated in Fig. 3B) area and size characteristics over time (z-score normalised), with corresponding p-values. Note, z-score in blue-to-red scale is size. The corresponding p-values are presented for these area changes wherein each condition compared to the value for the sgSCR (-CTX) as control. (**D)** Scoring of organised lumen-containing spheroids (as indicated in Fig. 3B) versus disorganised aggregates at 96 h (± cholera toxin), represented as fold change to control. Datapoint shapes correspond to each of 3 independent experiments. **E,** Immunofluorescence of HT-29 spheroids stained for PODXL, NHERF1/EBP50, F-actin and nuclei (SCR and PODXL KO ± CTX). PODXL immunolabelling pseudocoloured based on intensity, using the blue-to-red colour-mapping indicated. **F,** Schema, effect of cholera toxin (CTX) and PODXL KO on lumen formation and multicellular morphogenesis in HT-29 cell spheroids. *Statistical analyses: Student’s t-test. Note that sgPODXL 2 is used where written as sgPODXL, unless otherwise indicated*.

Given PODXL’s established function in epithelial lumen formation^14–16,41^, we hypothesised that it might instead control the collective organisation of CRC cells. To test this, we grew cells as 3D spheroids in ECM and monitored their development over several days by label-free time-lapse imaging (time course, 0–144 h), coupled to quantification of area as a proxy for growth. Control (SCR) HT-29 spheroids typically formed compact structures that, upon treatment with cholera toxin (CTX) - a known inducer of lumen formation in multiple cell types^49–51^ - developed multiple small internal lumens (Fig. 3B, black arrowheads). Consistent with PODXL having a minimal effect on 2-D growth, PODXL KO did not significantly alter overall spheroid size, irrespective of CTX (Fig. 3C, Area), but it profoundly altered spheroid organisation: PODXL KO spheroids showed an increased propensity to form disordered, grape-like aggregates lacking lumens, rather than cohesive, lumen-containing structures (Fig. 3B – image examples, Fig. 3D quantitation).

By immunofluorescence, CTX-treated control spheroids (SCR) showed PODXL at luminal surfaces (arrowheads) together with its binding partner NHERF1, and organised filamentous actin at cell junctions (Fig. 3E, left panel). In contrast, PODXL KO spheroids lacked lumens and luminal NHERF1 labelling, instead showing disorganised packing of neighbouring cells and reduced junctional filamentous actin (Fig. 3E, right panel). Together, these data are consistent with a role for PODXL in coordinating lumen-forming collective cell behaviour through the actin cytoskeleton (Fig. 3F).

To test this in a second, independent system, we depleted *PODXL* by shRNA in KPN liver-metastasis-derived organoids (KPN-Lmet1; Fig. 4A) and again performed time-lapse imaging. The resulting 3D structures were smaller than controls, in a knockdown-efficiency-dependent manner (Fig. 4B–D). More strikingly, and mirroring effects seen in HT-29 cells, PODXL depletion caused a shift away from spherical, lumen-containing organoids toward disorganised cell aggregate structures (Fig. 4B,E). This was not a consequence of lack of cell density, as this phenotype was maintained at two different plating densities (Fig. 4D; 1.5 or 2.4 x10^4^ cells/ml). As PODXL KO HT-29 cells and KD murine organoids were not clonally selected following CRISPR editing, it remains possible that the minority of spheroids able to form organised structures arose from residual PODXL-expressing cells within the population.

**Figure 4.**
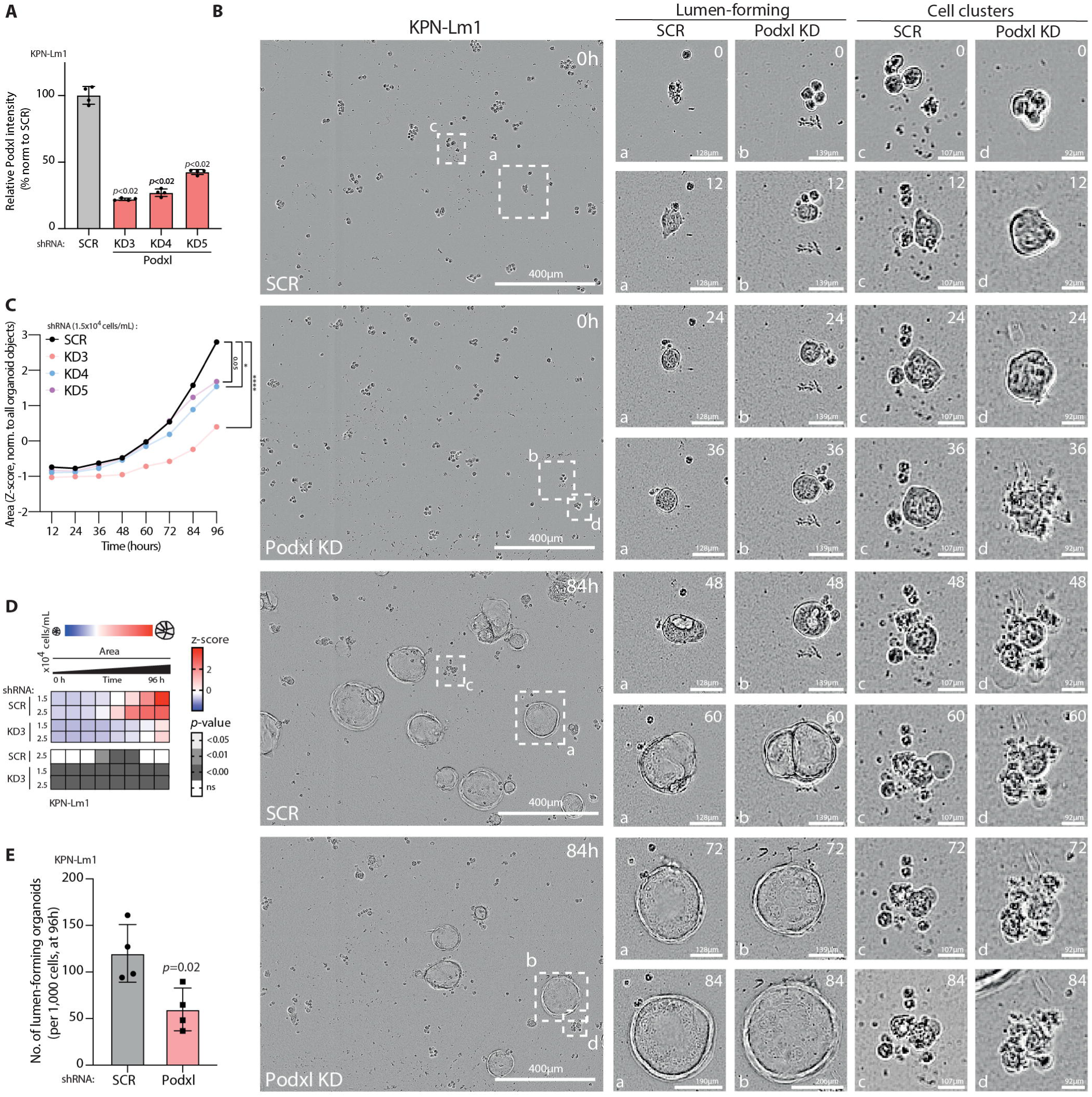
PODXL knockdown disrupts collective organisation and lumen formation in CMS4-like KPN organoids. **A,** qPCR confirmation of PODXL knockdown in KPN liver-metastasis–derived (Lm1) organoids (shRNA scramble, KD3–KD5). n = 4 independent experiments. **B,** Representative phase-contrast images from one experiment of live Incucyte imaging of organoid formation across 4 independent experiments (each with two alternative plating densities. These representative images come from cells plated at 1.5 x 10^4^ cells/ml) with 4 technical replicates in each; left, field views; right, time courses of individual lumen-forming organoids and disorganised cell clusters. **C-D,** Analyses if all experimental images corresponding to Fig. 3B, not only images shown. **(C)** Organoid area over time (z-score normalised) for scramble and *Podxl* shRNAs; statistical analysis at 96 h (calculated across all experiments indicated in Fig. 4B). **(D)** Heatmap of organoid size characteristics across two different plating densities (1.5 x 10^4^ or 2.5 x 10^4^ cells/ml) and time (z-score normalised) calculated across all experiments indicated in Fig. 4B. Note, Z-score in blue-to-red scale is size. The corresponding p-values are presented for these area changes wherein each condition compared to the value for the KPN-Lm1 scramble line (at 1.5 x10^4^cells/ml) as control. **E,** Organoid formation at 96 h (number of organoids per 1,000 cells; scramble vs *Podxl* shRNA). Datapoint shapes correspond to each of 4 biological replicates. *Statistical analyses: Mann–Whitney test (A), one-way ANOVA (C, D), Student’s t-test (E)*.

Nevertheless, the consistent shift toward disorganised morphologies across independent human and murine 3-D systems supports a requirement for PODXL in collective morphogenesis. Together, these findings demonstrate that PODXL promotes the organisation of CRC cells into cohesive, lumen-containing structures in 3-D culture.

### PODXL promotes tumour growth and glandular architecture *in vivo*

To determine whether PODXL’s role in collective organisation extends to tumours in vivo, we transplanted control (SCR) and PODXL KO HT-29 cells into immunocompromised mice. We first assessed subcutaneous tumour formation, which allows straightforward monitoring of tumour growth (Fig. 5A). Both control and PODXL KO HT-29 cells formed tumours with 100% efficiency (Fig. 5B). Quantitative immunohistochemical analysis confirmed a sustained reduction in PODXL levels in KO tumours (Fig. 5C). One PODXL KO tumour completely regressed and was therefore unavailable for analysis, resulting in an *n* = 7 for the KO group. In control tumours (SCR), PODXL protein was localised to glandular lumens (Fig. 5D, arrowheads), mirroring its pattern in the patient TMA (Fig. 1I) and GEMM tumours (Fig. 2D), and consistent with a conserved function in lumen organisation. Although initial tumour establishment was unaffected by attenuated PODXL expression, tumour growth diverged over time. Control tumours (SCR) increased steadily in volume over the five-week experiment, whereas PODXL KO tumours plateaued after approximately two weeks. Two PODXL KO tumours behaved as outliers, reaching the clinical endpoint during weeks 2 and 3 (pink points, Fig. 5E). The remaining PODXL KO tumours were significantly smaller than controls at weeks 4 and 5 (Fig. 5E). Mirroring effects in 2D and 3D growth ex vivo, this growth difference was not explained by changes in proliferation, as Ki67-positive cells were equivalent between groups (Fig. 5F), nor by increased cell death, as necrotic areas were similar (Fig. 5G). Instead, mirroring effects seen in 3-D culture of HT-29 (Fig. 3) and KPN organoids (Fig. 4), PODXL KO tumours showed a marked loss of glandular architecture (Fig. 5H), as shown by representative haematoxylin and eosin (H&E) staining (Fig. 5I). Control (SCR) tumours formed organised glandular structures with discernible lumens and mucin-containing vacuoles (Fig. 5I, left, arrowheads) - the two features by which glandular architecture was scored by a clinical pathologist (K.R.) - whereas PODXL KO tumours presented as solid, disorganised masses lacking luminal organisation (Fig. 5I, right panels). These findings indicate that PODXL does not primarily regulate tumour establishment, proliferation, or survival, but instead governs tissue-level organisation into glandular structures, which is necessary for efficient tumour growth.

**Figure 5.**
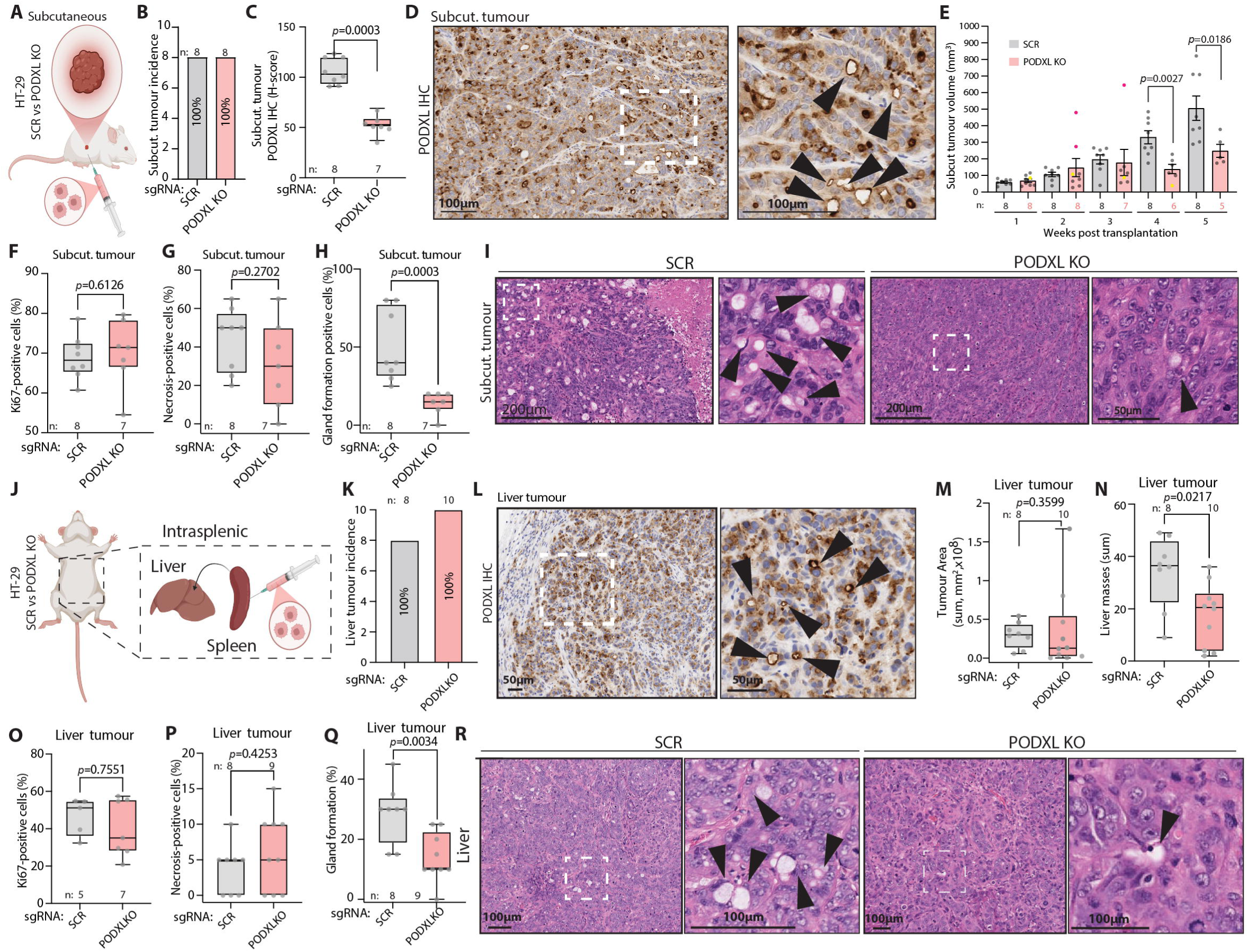
PODXL promotes tumour glandular architecture and metastatic colonisation in vivo. **A–I,** Subcutaneous xenograft model: schematic **(A)**, tumour incidence **(B)**, PODXL IHC quantification (Histoscore) **(C)**, representative PODXL IHC in SCR, apical/luminal labelling (arrowheads) **(D)**, tumour volume over time **(E)**, Ki67 quantification **(F)**, necrosis quantification **(G)**, gland-formation quantification **(H)**, and representative H&E images. Arrowheads indicated glands, with less present in PODXL KO **(I)**. In **(E)**, pink datapoints denote two “outgrower” PODXL KO tumours that reached clinical (humane) endpoint in weeks 2 and 3, before the timed endpoint of 5 weeks; the yellow point denotes one PODXL KO tumour that regressed to undetectable and was therefore not harvested or used for downstream analysis. Because these two outgrowers reached endpoint and were censored from subsequent timepoints, the reduced volume of PODXL KO tumours at later timepoints reflects the remaining, non-outgrower tumours. *Tumour volume was compared between groups at each timepoint by two-tailed Mann–Whitney test; PODXL KO tumours were significantly smaller than SCR controls at week 4 (p=0.0027) and week 5 (p=0.0186)*. **J-R** Intrasplenic xenograft model: **(J)** schematic, splenic vein draining to the liver, **(K)** liver metastasis incidence, **(L)** representative PODXL IHC of liver metastases (SCR), apical/luminal labelling (arrowheads), **(M)** liver tumour area, **(N)** metastatic masses in the liver (sum), **(O)** Ki67-positive cells (%) note, alternative sample number as pilot cohort was not scored, **(P)** necrosis-positive cells (%), **(Q)** gland-formation phenotype (%), **(R)** representative H&E images (SCR / PODXL KO); arrowheads indicate glandular structures. For liver metastases, tumour area (M) and metastatic mass number (N) were quantified for all mice (n=8 SCR / 10 PODXL KO); one PODXL KO liver metastasis was too small for reliable morphological assessment and, on the pathologist’s recommendation, was excluded from necrosis (P) and gland-formation (Q) scoring, giving n=8 SCR / 9 PODXL KO for those two panels. Statistical analysis: pairwise SCR vs PODXL KO comparisons (C, F, G, H, M, N, O, P, Q) were performed using a two-tailed Mann–Whitney U test; subcutaneous tumour volume (E) was compared between groups at each timepoint by two-tailed Mann–Whitney test. Tumour incidence (B, K) was 100% in the relevant groups and was not tested. *Box-and-whisker plots show median, interquartile range and min–max with individual mice as points; volume data are mean ± SEM. Exact p-values are shown on each panel. Scale bars indicated*.

### PODXL controls glandular architecture and metastatic burden in the liver

To determine whether PODXL contribution to tumour organisation extends to metastatic sites, we performed intrasplenic injections of control (SCR) and PODXL KO HT-29 cells (Fig. 5J). This model allows assessment of hepatic metastasis, as the splenic vein drains directly to the liver^52^. For liver metastasis, tumour incidence was 100% in both groups (Fig. 5K) (n = 8 SCR, 10 PODXL KO; two SCR excluded, see Methods), with PODXL once again localised to luminal structures in SCR liver metastases (Fig. 5L, black arrowheads). Although total tumour area was not significantly different (Fig. 5M), PODXL KO livers contained significantly fewer metastatic masses (Fig. 5N). As discrete lesions can be difficult to enumerate on two-dimensional sections, particularly where metastases are closely apposed or coalescing, we interpreted metastatic mass number alongside total tumour area (Fig. 5M) as complementary measures of metastatic burden. This reduction in metastatic lesion number, despite equivalent incidence and overall burden, suggests that PODXL KO cells can seed the liver but are impaired in establishing or expanding individual metastatic colonies. As with subcutaneous tumours, liver metastases showed no difference in proliferation (Fig. 5O) or necrosis (Fig. 5P) but displayed dramatically reduced gland formation (Fig. 5Q). Representative histology showed that control (SCR) liver metastases formed organised glandular structures (Fig. 5R, left panels, arrowheads), while KO metastases appeared as solid, disorganised masses (Fig. 5R, right panels). This recapitulates the loss of glandular organisation seen upon PODXL depletion in vitro and in subcutaneous tumours, indicating that PODXL supports the same lumen-forming programme at the metastatic site. Collectively, these data indicate that PODXL governs glandular tumour architecture across primary and metastatic sites, and influences metastatic burden in the liver, without affecting tumour initiation.

## Discussion

Tumours frequently co-opt developmental gene programmes to drive progression, yet the functional contribution of genes controlling tissue organisation has remained unclear. Our findings demonstrate that PODXL - a key regulator of apical-basal polarity and epithelial lumen formation during development^14,16,17,26^ - promotes glandular architecture in CRC. This organised structure actively contributes to tumour efficiency and metastatic colonisation. Rather than representing a vestigial feature that tumours progressively lose, glandular organisation enabled by PODXL confers functional advantages within tumours.

PODXL’s contribution to CRC became apparent only in contexts that permitted tissue-level organisation. In HT-29 cells, PODXL depletion produced no detectable phenotype in standard 2D culture; effects emerged exclusively in 3D settings, where PODXL loss disrupted collective organisation into lumen-containing structures without affecting spheroid size. This organisational defect was also conferred in vivo, with PODXL deficiency resulting in loss of glandular architecture - defined as the presence of both organised luminal structures and mucin-containing vacuoles - largely in the absence of changes in proliferation (Ki67) or necrosis. PODXL therefore governs tumour tissue organisation rather than tumour cell fitness and explains why PODXL would appear phenotypically dispensable in conventional proliferation-based/2-D assays. In CRC, preservation of glandular architecture is a defining feature of well-differentiated tumours and is generally associated with more favourable clinical outcomes, whereas the prognostic significance of mucinous differentiation is context-dependent and varies with tumour subtype and growth pattern^53^. Notably, mucinous histology is most frequently observed in CMS1 and CMS4 CRCs^43,54^.

Loss of glandular architecture had clear functional consequences. Despite comparable tumour cell seeding, PODXL-deficient subcutaneous tumours were, on average, smaller at later timepoints - although two PODXL KO tumours reached the humane endpoint (maximum permitted size) during weeks 2-3 and were censored from later timepoints - and generated significantly fewer liver metastases. Organised glandular architecture itself may therefore contribute to efficient tumour growth and metastatic colonisation. PODXL-deficient cells can seed and survive, but fail to organise efficiently, suggesting that glandular architecture provides structural advantages that support sustained tumour expansion. In breast cancer, maintenance of cell-cell adhesion through E-cadherin, and the formation of microlumens between cancer cells, is required for metastasis by regulating spatial signalling between neighbouring metastatic cells^3,37^. This is analogous to collections of cells that migrate during development, such as the developing lateral line cells in Zebrafish, that use microlumens between cells in a moving cluster to direct signalling within the cell cluster^55^. PODXL-dependent lumen formation may be a mechanism for controlling such tumour organisation.

The molecular basis of PODXL-mediated lumen formation in tumours likely parallels developmental mechanisms. In kidney epithelial cells, the PODXL/NHERF/Ezrin complex couples apical membrane identity to actin cytoskeletal organisation ^20,22,34,56^, and disruption of the PODXL-Ezrin interaction prevents lumen formation even when apical targeting is preserved^14^. PODXL-deficient CRC cells in our study exhibited reduced filamentous actin at cell-cell junctions, and previous work has shown that Ezrin inhibition reverses PODXL-dependent phenotypes in breast cancer^34^. Whether these same interactions mediate the broader glandular architecture phenotype - including mucin vacuole formation - will require structure-function analyses using PODXL domain mutants.

Our identification of TGF-β signalling as an upstream regulator of PODXL provides mechanistic insight into its selective elevation in CMS4, and to a lower extent CMS1, CRCs - a subtype defined by high TGF-β pathway activity, stromal infiltration, and poor prognosis ^43,57–60^. In CMS4 tumours, TGF-β released by stromal cells^60^ could drive epithelial PODXL expression in cells with intact signalling, even though some CRC cells can harbour inactivating mutations in canonical TGF-β pathway components^58,61^. It is interesting to note that CMS4 GEMM organoids, and human CRC cell lines had weaker correlation of TGFB3 expression to PODXL mRNA, suggesting some stromal regulation of TGF-β signalling to promote epithelial PODXL expression. This indicates that additional regulatory mechanisms maintain baseline PODXL expression while TGF-β drives elevation in aggressive tumours. Our functional analyses were performed using CMS4-like GEMMs (KP and KPN) and HT-29 xenografts; whether PODXL plays a similar architectural role in CMS2- or CMS3-like tumour contexts remains to be determined. Additionally, the high PODXL expression observed in CMS1 tumours - which in part may be attributed to immune and endothelial cell populations within the tumour microenvironment - warrants further investigation using single-cell or spatial transcriptomic approaches to deconvolve epithelial versus stromal contributions to PODXL expression across subtypes.

These findings have several clinical implications. First, the validation of PODXL as a prognostic marker specifically in CMS4 patients - at both mRNA and protein levels in independent cohorts - supports its potential utility for patient stratification. These prognostic associations were detected within a stage II–III, curatively resected cohort - the setting in which prognostic stratification is most clinically actionable. That a significant effect emerged even in this relatively homogeneous group suggests *PODXL*’s prognostic value may be greater still in cohorts spanning all stages, particularly those enriched for advanced or metastatic disease in which *PODXL*-driven glandular architecture and metastatic colonisation are most active. Notably, high PODXL protein was associated with reduced survival across all patients in the TMA analysis, whereas the mRNA-level association was restricted to CMS4. This discrepancy may reflect the weak association of PODXL protein and transcript level. Second, the demonstration that PODXL promotes tumour progression through effects on tissue organisation rather than proliferation suggests that targeting PODXL might complement conventional anti-proliferative therapies. Several groups have developed antibodies targeting the PODXL extracellular domain, which attenuate tumour formation and metastasis in experimental models^62,63^. Such approaches might be particularly effective in CMS4 CRC, where TGFβ–driven PODXL elevation supports organised tumour architecture.

In conclusion, this study identifies PODXL as a regulator of tumour glandular architecture in CRC, encompassing both lumen formation and mucin vacuole development. Rather than functioning primarily as a driver of individual cell metastasis, PODXL enables collective epithelial polarisation that supports organised tissue architecture and efficient tumour–host integration during metastatic colonisation. These findings support an emerging conceptual paradigm shift: tumours do not merely lose epithelial organisation during progression but may actively maintain glandular architecture to drive disease advancement. The co-option of developmental epithelial programmes represents a previously underappreciated mechanism by which tumour cells achieve efficient growth and metastatic colonisation.

## Methods

### Patient cohort analyses

#### TCGA data analysis

Publicly available colorectal cancer (CRC) datasets were analysed to assess associations between *PODXL* expression, molecular subtype, and clinical outcome. Primary analyses used the TCGA PanCancer Atlas (2018)^62^, including RNA-sequencing data from colon adenocarcinoma (COAD) and rectal adenocarcinoma (READ) with matched clinical annotations obtained via cBioPortal (www.cbioportal.org). Pan-cancer analyses were performed using the ICGC/TCGA Pan-Cancer Analysis of Whole Genomes (PCAWG) cohort (2020)^63^. *PODXL* mRNA expression was quantified as log_2_-transformed transcripts per million (TPM), and copy-number alteration data were retrieved from the same source.

#### Consensus Molecular Subtype classification

CMS classifications for TCGA and GEO datasets were obtained from the CRC Subtyping Consortium (CRCSC) Synapse repository (Synapse ID: syn2623706; https://www.synapse.org/Synapse:syn2623706/wiki/67246). These classifications represent the gold-standard consensus calls derived from the original CRCSC study, in which six independent subtyping algorithms were applied to harmonised gene expression data and network analysis was used to identify consensus clusters. Patients were classified into CMS1 (MSI Immune), CMS2 (Canonical), CMS3 (Metabolic), or CMS4 (Mesenchymal) subtypes. Samples that could not be confidently assigned to a single subtype were excluded from subtype-specific analyses.

#### Additional microarray datasets

Independent validation was performed using publicly available datasets from the Gene Expression Omnibus (GEO). Datasets using GPL96 (Affymetrix Human Genome U133A Array) and GPL570 (Affymetrix Human Genome U133 Plus 2.0 Array) platforms were analysed. Specific datasets indicated in figure panels and legends. For datasets containing laser capture microdissected samples, epithelial-enriched fractions were analysed separately.

#### Survival analyses

For survival analysis based on mRNA, patients were stratified by median split of *PODXL* mRNA expression into *PODXL*-high and *PODXL*-low groups. Overall survival was analysed using Kaplan-Meier curves and compared using log-rank tests. Analyses were performed both across all patients and within individual CMS subtypes. Hazard ratios were calculated using Cox proportional hazards regression.

### Tissue microarray analysis

#### Patient cohort

A retrospective cohort of 778 stage II-III CRC patients who underwent surgery with curative intent resection within Greater Glasgow and Clyde NHS (1997-2013) was analysed. Of these, 447 patients with scorable PODXL tissue microarray cores and Consensus Molecular Subtype (CMS) calls were included in the analyses reported here. Ethical approval was in place (MREC/01/0/36) and data are stored within the Glasgow Safehaven (GSH21ON009). Clinicopathological data including age, sex, tumour site, TNM stage, differentiation grade, and survival outcomes were obtained from medical records. CMS classification was performed using the CMScaller R package^64^ in RStudio.

#### Immunohistochemistry

All IHC was performed by the CRUK Scotland Institute Histology facility. Previously constructed tissue microarray (TMA) sections (4 μm) from formalin-fixed paraffin-embedded (FFPE) tissue were stained for PODXL (ab150358, Abcam) using a Leica Bond Rx autostainer. Sections were dewaxed (AR9222, Leica) and underwent antigen retrieval using ER2 solution (AR9640, Leica) for 30 minutes at 100°C.

Sections were rinsed in Bond Wash (AR9590, Leica) before peroxidase blocking (DS9263, Intense R kit, Leica, 5 minutes). After washing, the anti-PODXL antibody (ab150358, Abcam) was applied at 1/1,000 dilution for 40 minutes. After washing, rabbit EnVision secondary antibody (K4003, Agilent) was applied for 30 minutes, and staining was visualised using DAB and counterstained with haematoxylin, both solutions contained within the Intense R kit.

#### Scoring and analysis

*PODXL* expression was quantified using QuPath digital pathology software, version 0.4.3. A weighted histoscore was calculated considering both staining intensity (0-3) and percentage of positive cells, generating scores from 0-300. PODXL was scored only within the tumour-cell cytoplasm; stromal expression, present in some cases, was not quantified (K.A.F.P.). An optimal threshold for dichotomisation of the weighted histoscore was determined using maximally selected rank statistics based on the log-rank test, implemented using the survminer package in R (v3.4.2). Patients were subsequently stratified into high- and low-expression groups according to this data-driven cut-point. Cancer-specific survival was analysed using Kaplan–Meier curves and log-rank tests.

### Correlation analyses

Correlations between the expression of *PODXL* mRNA and TGF-β pathway genes (*TGFB1*, *TGFB2*, *TGFB3*, *TGFBR1*, *TGFBR2*) were calculated using Pearson’s correlation coefficient. Correlation matrices were generated across multiple datasets including TCGA patient tumours, Cancer Cell Line Encyclopaedia (CCLE) cell lines, GEMM tumours, and GEMM-derived organoids. Statistical significance was determined using two-tailed t-tests with Bonferroni correction for multiple comparisons.

### Mouse models

#### Genetically engineered mouse models

GEMMs in this study have been previously described^46^. Briefly, KP mice carry villin-CreERT2-driven intestinal-specific activation of oncogenic *Kras* (*Kras^G12D/+^*) and p53 deletion (*Trp53^fl/fl^*). KPN mice additionally carry a *Rosa26*-targeted *Notch1* intracellular domain transgene (*Rosa26^N1ICD/+^*) that confers CMS4-like molecular features and metastatic capacity. KPN *Alk5^-/-^*(KPNA5) mice carry an additional intestinal-specific deletion of TGF-β receptor 1 (*Tgfbr1*/*Alk5*). Tumour induction was achieved by tamoxifen administration. Primary tumours and liver metastases were harvested at clinical endpoint for analysis.

#### Organoid derivation and culture

Organoids were supplied by Sansom lab. Organoids were maintained in Complete Culture Medium (CCM): Advanced DMEM/F12 supplemented with 2 mM L-Glutamine, 10 mM HEPES, 1× B27, 1× N2, 50 ng/ml EGF, and 100 ng/ml Noggin. Organoids were passaged every 2-3 days by mechanical dissociation and re-embedding in fresh Growth Factor Reduced Matrigel (GFRM; BD Biosciences, 354230) in domes. For passaging, spent medium was removed, organoids were washed and scraped with cold PBS (∼1 mL per 20 μL dome), mechanically dissociated by vigorous pipetting, and centrifuged at 400 × g for 5 minutes at 4°C. After a second wash and centrifugation at 600 × g, organoids were resuspended in fresh GFRM at 1:1.5-3 split ratio and plated as 20 μL domes on 6-well plates (4-7 domes per well). Plates were inverted and incubated at 37°C for 10 minutes to allow polymerisation, then 2.5 mL CCM was added per well.

### Cell lines and culture

#### Cell line maintenance

HT-29 (female, ATCC HTB-38) human CRC cell line was maintained in DMEM supplemented with 10% FBS, 2 mM L-Glutamine, and 1% penicillin/streptomycin. Cells were cultured at 37°C with 5% CO2 and passaged every 2-4 days at approximately 80-90% confluence. Additional cell lines used were cultured as per ATCC recommendations. Cell lines were authenticated by STR profiling and confirmed mycoplasma negative.

#### Generation of PODXL knockout and knockdown lines

*PODXL* knockout was achieved using CRISPR-Cas9. sgRNA sequences targeting *PODXL* exons were cloned into pLentiCRISPR v2 vector (Addgene #52961) with puromycin resistance. sgRNA sequences used:

- sgPODXL-1: AATGCCGTTGCCGGGCTCGT
- sgPODXL-2: AGATAAGTGCGGCATACGGC
- Scramble control: GACCGGAACGATCTCGCGTA

For knockdown experiments in organoids, shRNA sequences targeting mouse *Podxl* were cloned into pLKO.1 vector:

- shPodxl-1: CTTTGGAAATTACCAGCTAAA
- shPodxl-2: GCTCTTCATTTCAGTGGCAAA
- Scramble control: CCGCAGGTATGCACGCGT

All sgRNA and shRNA constructs were designed against regions conserved across both *PODXL* isoforms, ensuring loss of expression of all annotated *PODXL* isoforms.

#### Lentiviral production and transduction

HEK293FT (female) packaging cells were seeded at 2.5 × 10^5^ cells per well in 6-well plates for 24 hours. Cells were co-transfected with lentiviral packaging vectors pMD2.G (VSVG; Addgene #12259) and psPAX2 (Addgene #12260) together with the plasmid of interest using Lipofectamine 2000. Specifically, 0.5 μg plasmid, 0.55 μg psPAX2, 0.05 μg VSVG, and 6 μL Lipofectamine 2000 were diluted in 500 μL Opti-MEM and added to cells. Medium was changed the following day to growth medium plus 10% additional FBS. Viral supernatants were collected at 48 and 72 hours post-transfection, clarified by centrifugation (500 × g, 5 minutes), filtered through 0.45 μm PES filters, and concentrated using Lenti-X Concentrator (Takara, 3:1 ratio, 4°C for 1 hour). Concentrated virus was collected by centrifugation (1,500 × g, 45 minutes, 4°C) and resuspended in PBS.

For cell line transduction, target cells were seeded at 1.5-3 × 10^5^ cells per well in 6-well plates. Concentrated virus was added in culture medium with 8 μg/mL polybrene. After 48 hours, cells were selected with appropriate antibiotics (puromycin 2 μg/mL or blasticidin 10 μg/mL).

#### TGF-β treatments

For TGF-β induction experiments, cells were serum-starved overnight then treated with recombinant human TGF-β1, TGF-β2, or TGF-β3 (R&D Systems) at 10 ng/mL for 24 hours. For TGF-β inhibition, cells were treated with SB431542 (Tocris, 1614) at 10 μM. For shRNA-mediated knockdown of TGFBR1, cells were transduced with pLKO.1 vectors containing the targeting sequences below and selected with puromycin. shRNA sequences used (human TGFBR1):

- shTGFBR1-1: GAAGTTGCTGTTAAGATATTC
- shTGFBR1-2: GATCATGATTACTGTCGATAA
- shTGFBR1-3: GCTGGTCTTAACTTTAGGTAA
- shTGFBR1-4: CTCATGTTGATGGTCTATATC
- shTGFBR1-5: CGATGTTCCATTGGTGGAATT

### Biochemistry

#### Cell lysate preparation

Cells at 80% confluence in 6-well plates were washed twice with ice-cold PBS and lysed in 150 μL RIPA buffer (50 mM Tris pH 7.4, 150 mM NaCl, 1% NP-40, 0.25% sodium deoxycholate) supplemented with protease inhibitor cocktail (Complete Ultra, Roche) and phosphatase inhibitors (PhosStop, Roche). Lysates were incubated on ice for 15 minutes and clarified by centrifugation at 13,000 × g for 15 minutes at 4°C.

#### Protein concentration assay

Protein concentration was determined using the Micro BCA Protein Assay kit (Thermo Fisher Scientific) according to the manufacturer’s instructions, with BSA standards.

#### Western blotting

Equal amounts of protein (20 μg) were resolved on Bolt 4-12% Bis-Tris Plus gels (Thermo Fisher Scientific) in MOPS buffer at 120V for 80 minutes. Proteins were transferred to PVDF membranes using the iBlot 2 system (7 minutes). Membranes were blocked in Rockland blocking buffer for 1 hour and incubated with primary antibodies overnight at 4°C: anti-PODXL (ab150358, Abcam, 1:1,000), anti-GAPDH (2118, Cell Signaling Technology, 1:5,000). After washing with TBS-T, membranes were incubated with fluorescent secondary antibodies (DyLight 800, 1:10,000) or HRP-conjugated secondary antibodies (1:5,000) for 1 hour at room temperature. Detection was performed using Li-Cor Odyssey CLx scanner (fluorescent) or Bio-Rad ChemiDoc system (chemiluminescent, SuperSignal West Pico/Femto).

### RNA extraction and quantitative PCR

#### RNA extraction

Total RNA was extracted using the RNeasy Mini Kit with QIAshredder homogenisation (Qiagen). Cells at 70% confluence were washed twice with PBS, lysed in 350 μL RLT buffer containing β-mercaptoethanol, and homogenised through QIAshredder columns. RNA was purified according to the manufacturer’s instructions and eluted in 50 μL nuclease-free water. RNA concentration and purity were measured using a NanoDrop spectrophotometer.

#### Reverse transcription and qPCR

cDNA was synthesised from 1.5 μg RNA using the High-Capacity cDNA Reverse Transcription kit (Applied Biosystems). qPCR was performed using PowerUp SYBR Green Master Mix (Thermo Fisher Scientific) on an Applied Biosystems QuantStudio 3 system. Primers were used at 2 μM per reaction. Cycling conditions: 50°C 2 minutes (UNG incubation), 95°C 10 minutes (polymerase activation), then 40 cycles of 95°C 15 seconds (denature) and 61.5°C 1 minute (anneal/extend), followed by melt curve analysis. Three independent RNA samples were prepared per condition with four technical replicates each. Relative expression was calculated using the ΔΔCt method with *GAPDH* as reference gene.

Primer sequences:

- Human *GAPDH* Forward: CCCTTAAGAGGGATGCTGCC
- Human *GAPDH* Reverse: TACGGCCAAATCCGTTCACA
- Human *PODXL* Forward: AGTGACATGAAGCTGGGGAA
- Human *PODXL* Reverse: ACCATTCTCCACTGTCTGCA

### Three-dimensional culture and imaging

#### 3-D culture method for live imaging

For 3-D culture, 96-well ImageLock plates (Sartorius) were pre-coated with 10 μL Growth Factor Reduced Matrigel per well and incubated at 37°C for 10 minutes. Cell lines were trypsinised, counted, and resuspended at 1.5 × 10^4^ cells/mL in culture medium supplemented with 2% Matrigel. 150 μL of cell suspension was added per well. For organoids, cultures were dissociated to small clusters using StemPro Accutase (5 minutes at 37°C), counted, and resuspended at 2 × 10^4^ cells/mL in CCM supplemented with 2% Matrigel. Plates were equilibrated at 37°C for 30 minutes before imaging. Media was refreshed every 2-3 days.

#### Cholera toxin treatment

For lumen induction experiments, cholera toxin (Sigma, C8052) was added at 0.05 μg/mL to 3-D cultures at the time of plating.

#### Live imaging

Live imaging was performed using an Incucyte S3 Live Cell Analysis System (Sartorius). Using scratch-wound imaging mode, 2 positions per well were captured to focus on the plate grid and avoid edge effects. Phase contrast images were acquired every hour for up to 7 days.

#### Image analysis

Exported image series were analysed using custom pipelines in CellProfiler and KNIME Analytics Platform^65^. Per-object measurements including area, perimeter, form factor, eccentricity, compactness, and aspect ratio were extracted. For morphological classification, objects were categorised as organised cysts (spherical, compact structures with or without lumens) or aggregates (irregular, grape-like clusters) based on form factor and compactness thresholds determined from training datasets. Machine learning-derived classification was performed using Random Forest classifiers trained on manually annotated datasets.

#### Immunofluorescence of 3-D structures

3-D cultures were fixed with 4% paraformaldehyde for 15 minutes at room temperature, washed twice with PBS, and blocked in PFS buffer (0.7% fish skin gelatin in 0.025% saponin-PBS) for 1 hour with gentle agitation. Primary antibodies were diluted in PFS and incubated overnight at 4°C with gentle agitation: anti-PODXL (18150-1-AP, Proteintech, 1:100), anti-NHERF1/EBP50 (ab9526, Abcam, 1:100). After three 5-minute washes with PFS, secondary antibodies (Alexa Fluor 488 or 647 conjugated, 1:200), Alexa Fluor 568-phalloidin (1:200), and Hoechst 34580 (1:1,000) were added for 1 hour at room temperature. Following three washes with PFS and three washes with PBS, plates were imaged.

#### Confocal imaging

Fixed and stained 3-D cultures were imaged on an Opera Phenix High Content Screening System (PerkinElmer) using a 63× water immersion objective. Z-stacks were acquired at 2 μm intervals. Maximum intensity projections were generated, and images were analysed using Columbus software (PerkinElmer) with custom analysis pipelines.

### Xenograft studies

#### Animal husbandry

All animal experiments were carried out in compliance with UK Animals (Scientific Procedures) Act 1986 and EU directive 2010, under Home Office Project Licence PP6345023. Approval was obtained from the local Animal Welfare and Ethical Review Board (AWERB), University of Glasgow. Mice were kept in individually ventilated cages (IVCs) with controlled temperature (19-22°C) and humidity (55 ± 10%), 12-hour light/dark cycle, with access to food and water ad libitum, plus environmental enrichment.

#### Subcutaneous xenografts

Ten-week-old female CD-1 nude mice were randomly assigned to experimental groups and injected subcutaneously in the flank with 1.5 × 10^6^ HT-29 cells (Scramble control or *PODXL* knockout) in 100 μL PBS. Tumours were measured three times weekly using digital callipers by technical staff blinded to experimental group, and tumour volume was calculated as (length × width^2)/2. Mice were culled at humane endpoint (tumour diameter 15 mm) or at experimental endpoint (5 weeks). n = 8-13 mice per group across independent experiments.

#### Intrasplenic injection

Intrasplenic transplantation of HT-29 cells was run as a 3 SCR × 3 PODXL KO pilot experiment followed by a 7 SCR × 7 PODXL KO cohort, running in both instances to a timed end point of 6 weeks. In the 7×7 cohort, one SCR mouse was excluded prior to injection due to being a runt, and a second SCR mouse was culled 1 day post injection due to complications, resulting in final combined cohort sizes of 8 SCR and 10 PODXL KO mice. Female BALB/c nude mice (10 weeks old at pilot cohort; 10.8 weeks, 7×7 cohort) were randomly assigned to experimental groups and received analgesia (Carprofen, 5 mg/kg in drinking water) 24 hours before surgery and for 2 days post-surgery. Under general anaesthesia with Sevoflurane (7% induction, 3-5% maintenance), with additional Buprenorphine (0.05 mg/kg) and Carprofen (4 mg/kg) analgesia, mice were injected intrasplenically with 5 × 10^6^ HT-29 cells in 50 μL PBS. Mice were monitored for general health until endpoint (6 weeks post-implantation). At necropsy, livers and spleens were weighed, and tumour burden was assessed by assessors blinded to experimental group.

#### Histopathological analysis

Tissues were fixed in 10% neutral buffered formalin, processed, and embedded in paraffin. Sections (4 μm) were stained with haematoxylin and eosin (H&E) or processed for immunohistochemistry.

#### Immunohistochemistry of xenograft tissues

IHC was performed by CRUK Scotland Institute Histology facility. IHC was performed on a Leica Bond Rx autostainer. For PODXL staining, sections underwent antigen retrieval with ER2 solution (Leica) for 20 minutes at 95°C. After peroxidase blocking, anti-PODXL antibody (ab150358, Abcam) was applied at 1:1,000 dilution. Rabbit EnVision secondary (Agilent, K4003) was applied for 30 minutes, and staining was visualised with DAB. For Ki67, standard protocols were followed. Sections were counterstained with haematoxylin and coverslipped with DPX mountant.

#### Histological scoring

Macroscopic tumour measurements - subcutaneous tumour volume, and liver tumour area and metastatic mass number - were performed by L.G. Ki67, necrosis and gland-formation scoring was performed by a clinical pathologist (K.R), blinded to experimental group.

At dissection the liver was removed and separated into lobes and processed and stained as described. Haematoxylin and eosin (H&E) slides were scanned using an Aperio AT*2* slide scanner by the CRUK Scotland Institute Histology Facility and images uploaded to Halo v.3.6.4134.396 image analysis software from Indica. Using the annotation function, a layer was created to identify and define the boundary of each metastatic lesion within the total liver tissue on each slide. Using this layer, the software quantified the number of metastatic regions identified as well as the area of each lesion which summed gave the total area covered by all lesions in each section.

Necrosis and glandular differentiation were assessed visually on H&E sections and scored by a clinical pathologist (K.R.). Gland formation was scored as the proportion of viable tumour epithelium (across all lesions) forming organised, lumen-containing glands and/or mucin-containing vacuoles. Necrosis was scored as the proportion of total tumour area that was necrotic. Metastatic growth patterns were also assessed visually, described as either replacement (where the metastatic population fills the hepatocyte cell plate without inducing a stromal reaction) or desmoplastic (where there is induction of a stromal reaction of any sort). Both growth patterns were quantified in 5% increments. All visual histological scoring was performed by assessors blinded to experimental group.

For digital image analysis in QuPath (v0.4.3), a pixel classifier was built on Ki67-stained slides. Pixels were classified into tumour, stroma, immune and necrosis regions prior to running cell detection, using the random trees classifier at moderate resolution. Positive cell detection was then run within tumour areas, classifying them into ‘positive’ and ‘negative’.

### Statistical analysis

Tumour analyses were performed on n=8 SCR and n=7 PODXL KO subcutaneous tumours (one PODXL KO tumour regressed and yielded no tissue) and on the intrasplenic-cohort livers. One PODXL KO liver metastasis was too small for reliable morphological assessment and was excluded from necrosis and gland-formation scoring on the pathologist’s recommendation (n=8 SCR / 9 PODXL KO for those measures; it was retained for tumour-area and mass-count quantification).

## Data and code availability

TCGA data were obtained from cBioPortal (www.cbioportal.org). GEO microarray datasets are available at www.ncbi.nlm.nih.gov/geo/. CCLE data were obtained from the DepMap portal (depmap.org). Source data for all figures are provided with this paper. Custom analysis scripts are available from the first/corresponding author upon request.

## Acknowledgements

The authors acknowledge the Biological Services Unit, Flow Cytometry Facility (RRID:SCR_028243), Molecular Technology Services (RRID:SCR_027368), Histology Facility, Transgenic Models of Cancer, and the Beatson Advanced Imaging Resource (BAIR; Research Resource Identifier RRID: SCR_023875) at the CRUK Scotland Institute (RRID:SCR_027384 and ROR 03pv69j64). We acknowledge the BAIR team, in particular N. Paul and P. Thomason, for valuable discussions. We thank the Bryant laboratory for feedback, particularly E. Freckmann for support with image analysis and A. Roman-Fernandez for discussions regarding Podocalyxin. We thank G. Inman (CRUK Scotland Institute) for discussions regarding TGFβ signalling. We thank R. Ridgeway (CRUK Scotland Institute) for discussions regarding in vivo models. CRC GEMM and organoid RNA-seq data, and CRC organoids were provided by O. J. Sansom laboratory (CRUK Scotland Institute).

Illustrations were created using Adobe Illustrator and BioRender.com.

We are grateful to the patients whose samples enabled this research and acknowledge the use of murine models in this study. We thank C. Winchester (CRUK Scotland Institute) for critical review of the manuscript.

## Author Contributions

**E.M.C.**: Conceptualization, Methodology, Investigation, Formal Analysis, Visualization, Writing – Original Draft, Writing – Review & Editing. **K.R**.: Investigation (histological analysis), Formal Analysis. **K.A.F.P.**: Investigation (tissue microarray analysis), Formal Analysis. **L.A.G.**: Investigation (xenograft experiments). **E.S.**: Investigation (immunofluorescence), Writing – Review & Editing. **L.McG.**: Methodology (Operetta imaging). **R.J.**: Resources (organoids), Methodology. **C.N.**: Investigation (histology and immunohistochemistry). **L.M** and **K.B.**: Methodology, Supervision (xenograft studies). **J.LQ**: Supervision (histolopathological characterisation) **J.E.**: Supervision (histology and immunohistochemistry). **K.G.:** Data processing.**O.J.S.**: Resources, Methodology (GEMM expertise). **D.M.B.**: Conceptualization, Supervision, Funding Acquisition, Writing – Original Draft, Writing – Review & Editing.

## Supplementary info

### Mouse genotypes used in study

*All alleles are activated by intestinal-specific villin-CreERT2 following tamoxifen induction ^46^. Gene symbols are italicised; alleles are shown as superscripts*.

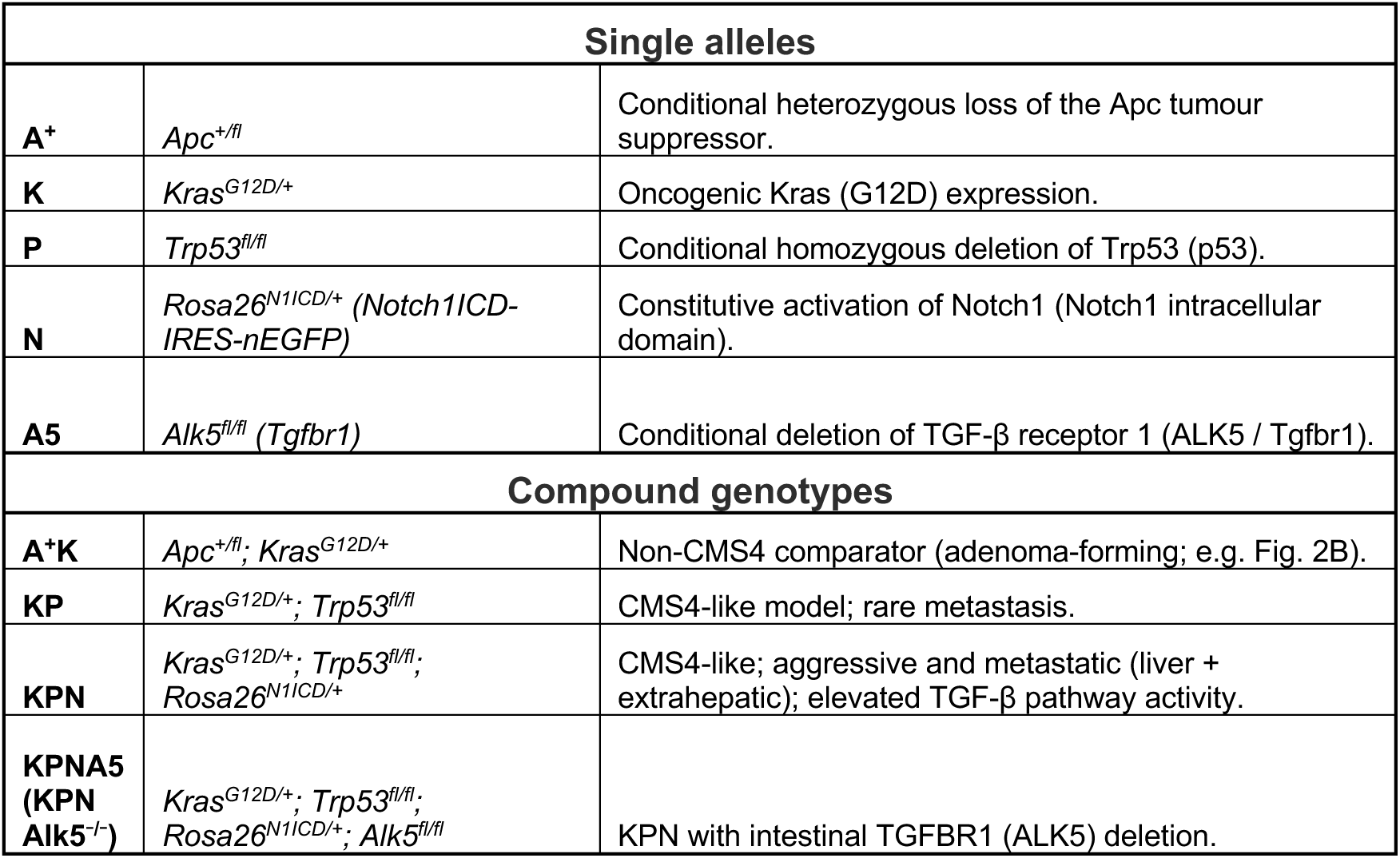

**Figure S1. Additional validation of *PODXL* mRNA expression across CRC datasets.**

**A–J,** *PODX*L mRNA in normal versus tumour tissue across the indicated TCGA (A-D) or GEO (E-J) datasets. Note that **C-D** are matched normal-tumour data (estimation plots).

**K–N,** *PODX*L mRNA in matched normal–tumour pairs across the indicated datasets (estimation plots).

**O,** Correlation between *PODXL* mRNA expression and copy number alteration in TCGA COAD+READ samples. Datapoint colour indicates copy status: Amplification, red; Gain, pink; Diploid, grey; Shallow deletion, blue.

**P–T,** *PODXL* mRNA expression in normal tissue, primary tumours, and metastases across indicated GEO microarray cohorts, analysed as unmatched (S) or matched (P, Q, R, T) samples.

**U-X,** *PODXL* mRNA stratified by tumour–node–metastasis (TNM) stage across the indicated cohorts.

**Y-Z,** *PODXL* mRNA expression in normal tissue, adenoma, and carcinoma across indicated microarray cohorts: unmatched (Y) and matched (Z) comparisons.

**AA–AF,** *PODXL* mRNA across CMS subtypes in the indicated independent cohorts.

*Box-and-whisker plots, Mann–Whitney; estimation plots, paired t-test. Patient sample numbers indicated*.

**Figure S2. PODXL expression association with patient outcome across CMS subtypes, sidedness and stage.**

**A,** Overall survival stratified by median-based *PODXL* mRNA, all patients, TCGA COAD + READ. **B,** Overall survival in CMS2; **C,** CMS3; **D,** CMS1 patients (*PODXL* mRNA), TCGA COAD + READ. **E,** *PODX*L mRNA by tumour sidedness (left vs right), TCGA COAD + READ.

**F,** Tumour (T) stage distribution stratified by *PODXL* mRNA, TCGA COAD + READ (stacked bar).

**G,** Cancer-specific survival in the CRC TMA cohort, all patients (PODXL protein, median split).

**H-K,** Survival from based on PODXL high versus low (optimised cutoff) expression in **(H)** Klintrup–Mäkinen (KM)-low or (**I),** Klintrup–Mäkinen (KM)-high tumours, **(J)** tumour–stroma-percentage (TSP)-low tumours, **(K)** CMS1, (**L),** CMS2; (**M,)** CMS3 or (**N),** CMS1+4 patients.

*Kaplan–Meier curves, log-rank (Mantel–Cox); box plots, Mann–Whitney; stacked bar, patient sample numbers indicated*.

**Figure S3. PODXL dependency and expression across colorectal cancer cell lines.**

**A,** Multi-metric heatmap of CRC cell lines: CMS classification, tumour origin (Primary versus Metastasis), *PODXL* mRNA expression, DepMap CRISPR dependency score, and mutation status of genes mutated in >10% of TCGA COAD/READ tumours.

**B, C,** *PODXL* mRNA expression when stratified by **(B)** CMS subtype or **(C)** primary versus metastasis-derived CRC cell lines.

**D,** Correlation between *PODXL* mRNA and DepMap dependency score; colours, CMS subtype.

**E, F,** PODXL dependency scores stratified by **(E)** CMS subtype or **(F)** tumour origin (primary versus metastasis-derived) for CRC cell lines.

**G,** Representative immunoblot of PODXL across CRC cell lines with GAPDH loading control; PODXL mRNA (log_2_ Transcripts per million, TPM) shown for comparison.

**H,** Quantification of total PODXL protein intensity normalised to the mean across 3 independent experiments, normalised to GAPDH. Alternative data point shapes correspond to each replicate.

**I,** Fraction of total PODXL protein per molecular-weight species from independent repeat experiments from H.

**J,** Correlation between PODXL protein abundance and *PODXL* mRNA in different cell lines.

**K-M,** Expression levels in HT-29 cells, when stably expressing either control (SCR, scramble) or each of five independent TGFBR1 shRNAs, for **(K)** *TGFBR1* mRNA (qPCR, % of SCR control), **(L)** *PODX*L mRNA (qPCR, fold change to scramble), **(M)** PODXL protein (fold change to scramble, normalised to GAPDH). A representative western blot is provided for PODXL and GAPDH. Alternative data point shapes correspond to 3 independent experimental replicates.

*Statistical analyses: Pearson correlation (D, J). n.s., not significant; sample numbers indicated*.

**Figure S4. PODXL knockout does not alter HT-29 2D colony formation or growth.**

**A-C,** In HT-29 cells expressing control (Scramble, SCR) versus PODXL KO **(A)** representative colony-formation images, and Superplot graphs of **(B)** colony count or **(C)** colony size from three independent experiments (large dots, smaller dots are the within experiment replicates).

**D,** Two-dimensional growth measured by number of nuclei, measured as nuclear integrated intensity over time (scramble vs PODXL KO), from two independent experiments in HT-29 cells expressing control (Scramble, SCR) versus PODXL KO.

*Statistical analyses: Mann-Whitney U-test*.

**Figure S5. PODXL loss in spleen tumours.**

Comparison of control (Scramble, SCR) and PODXL knockout (PODXL KO) HT-29 intrasplenic spleen primary tumours, combining a pilot (3 SCR × 3 PODXL KO) and a main (5 SCR × 7 PODXL KO) cohort. **A,** Spleen tumour incidence (SCR 8/8, 100%; PODXL KO 9/10, 90%).

**B,** Representative PODXL IHC; arrowheads indicate apical/luminal PODXL labelling; scale bar, 50 µm.

**C,** Number of tumour masses per spleen (n = 8 SCR / 10 PODXL KO).

**D,** Total tumour area per spleen (n = 8 / 10).

**E,** Ki67-positive tumour cells (%) (n = 5 / 7), note, alternative sample number as pilot cohort was not scored.

**F,** Necrosis (% of tumour) (n = 8 / 9).

**G,** Gland formation (% of tumour) (n = 8 / 9).

**H,** Representative H&E images (SCR / PODXL KO); arrowheads indicate glandular structures; scale bars, 50 µm.

Macroscopic tumour measurements (incidence, mass number and area; A, C, D) were performed by

L.G. using Halo image analysis; Ki67 (E; QuPath), necrosis (F) and gland formation (G) were scored by a clinical pathologist (K.R.). One PODXL KO mouse (YX6.7s) formed no tumour and is included as zero for mass number (C) and area (D) but excluded from Ki67, necrosis and gland-formation scoring; Ki67 (E) was quantified for the main cohort only (n = 5 SCR / 7 PODXL KO).

*Data are box-and-whisker plots (median, interquartile range; whiskers, minimum to maximum). Statistical analysis: SCR versus PODXL KO by two-tailed Mann–Whitney test. (C) p = 0.4982; (D) p = 0.5726; (E) p = 0.0101; (F) p = 0.755; (G) p = 0.1675*.

