## Supplementary figures and images for "Glandular architecture and malignant behaviour in colorectal cancer is regulated by the sialomucin Podocalyxin"

### Fig S1

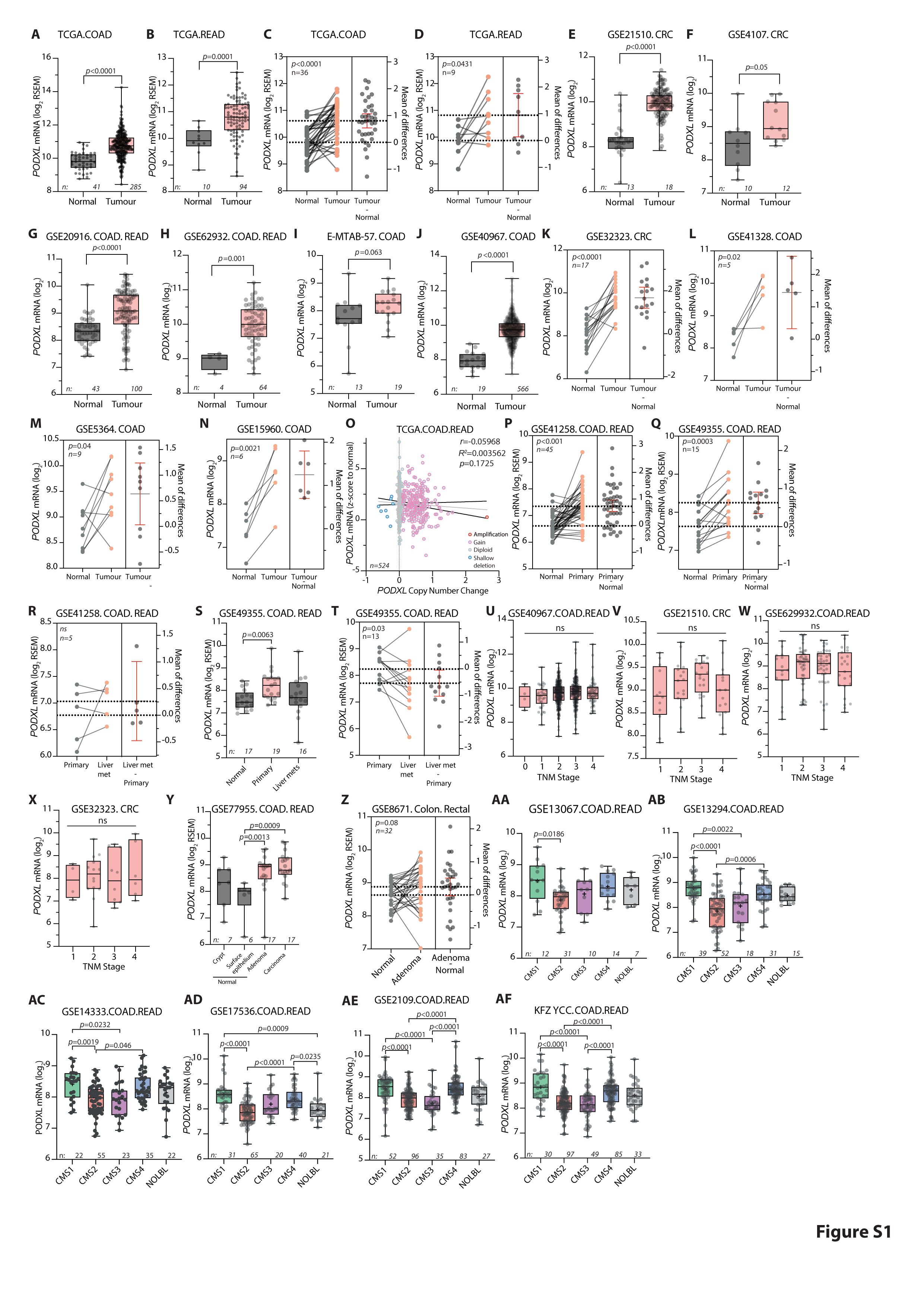

### Fig S2

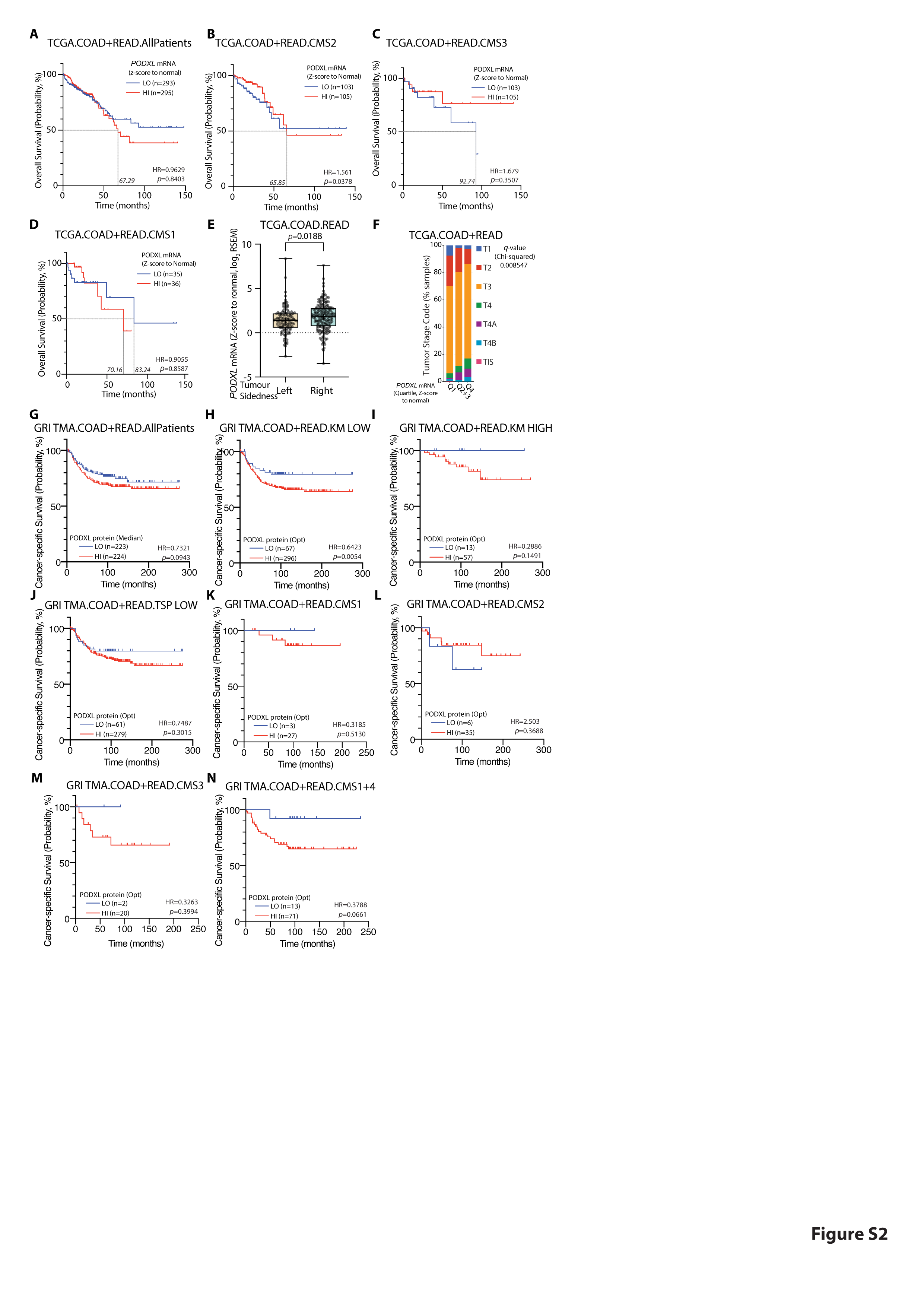

### Fig S3

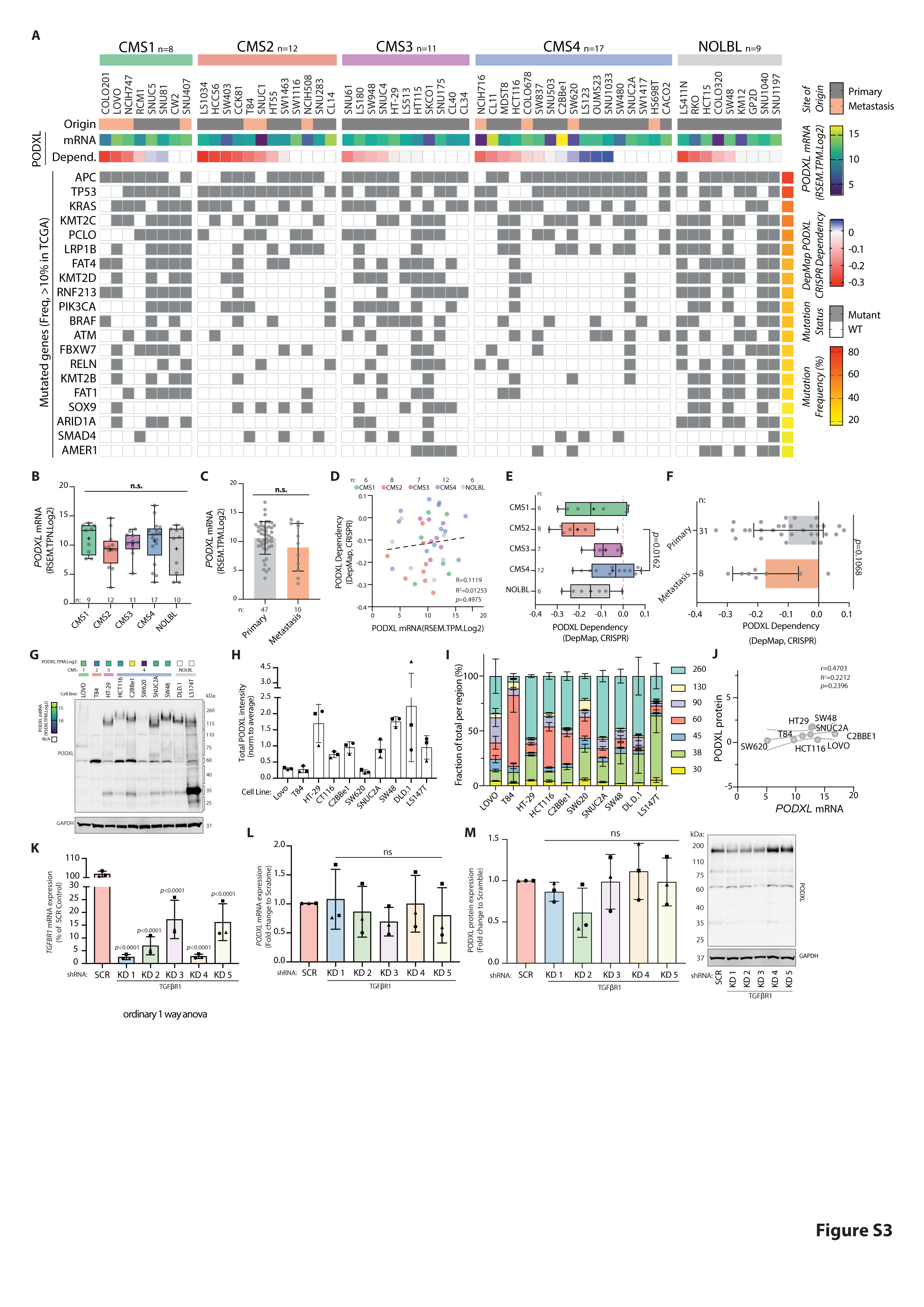

### Fig S4

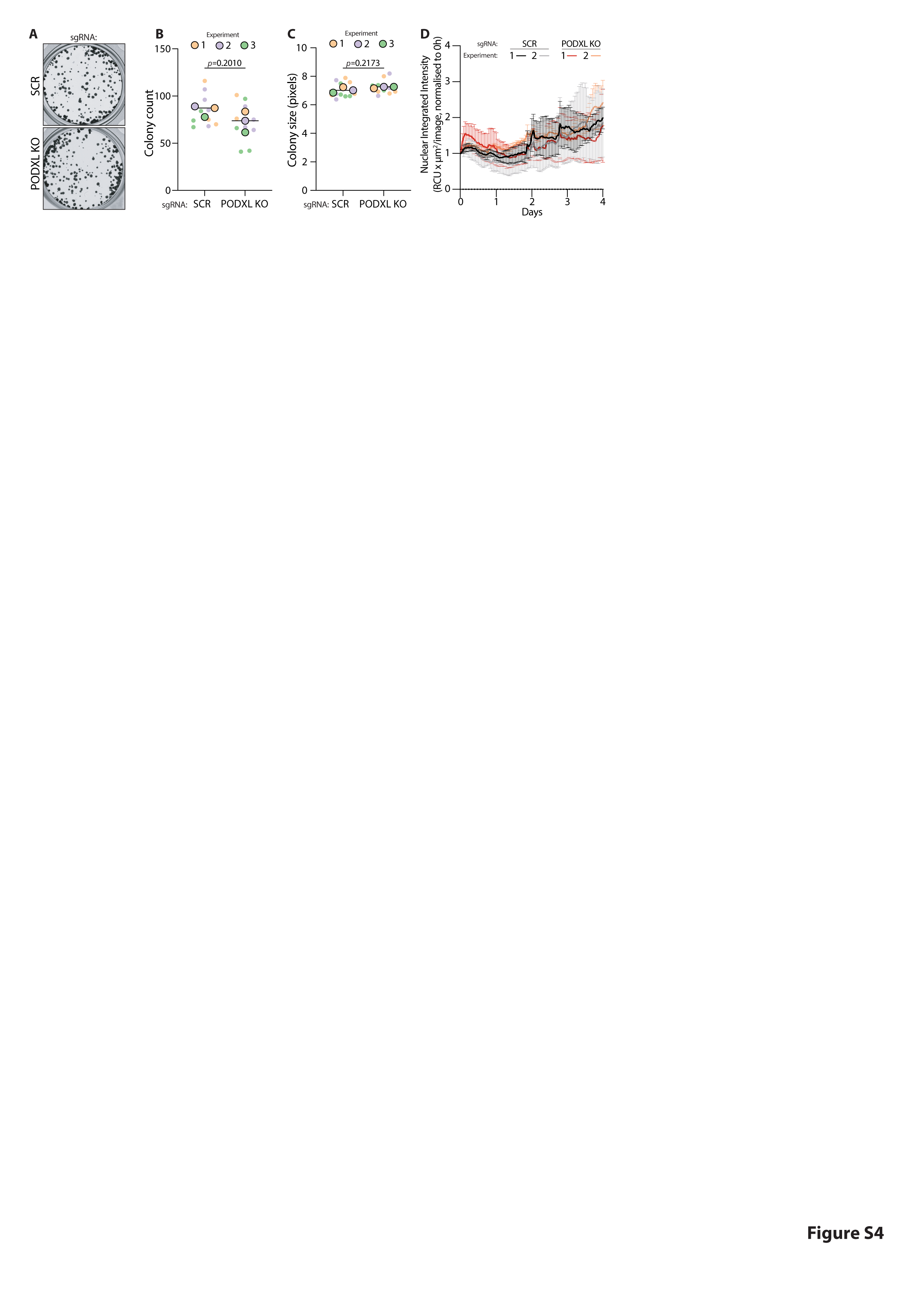
