## Supplementary material for "Glandular architecture and malignant behaviour in colorectal cancer is regulated by the sialomucin Podocalyxin": Table S1

| Clinical Feature | Low PODXL | High PODXL | p |
|---|---|---|---|
| **Sex** | | | |
| Female | 109 (47.4) | 102 (47.0) | 0.935 |
| Male | 121 (52.6) | 115 (53.0) | |
| **Age** | | | |
| <65 | 66 (28.7) | 65 (30.0) | 0.886 |
| 65-74 | 77 (33.5) | 68 (31.3) | |
| >75 | 87 (37.8) | 84 (38.7) | |
| **Tumour subsite** | | | |
| Right | 97 (42.2) | 106 (48.8) | 0.288 |
| Left | 72 (31.3) | 65 (30.0) | |
| Rectum | 61 (26.5) | 46 (21.2) | |
| **T stage** | | | |
| 1 | 15 (6.5) | 11 (5.1) | 0.071 |
| 2 | 32 (13.9) | 24 (11.1) | |
| 3 | 128 (55.7) | 113 (52.1) | |
| 4 | 55 (23.9) | 69 (31.8) | |
| **N stage** | | | |
| 0 | 144 (62.6) | 122 (56.2) | 0.213 |
| 1 | 57 (24.8) | 63 (29.0) | |
| 2 | 29 (12.6) | 32 (14.7) | |
| **Recurrence** | | | |
| Absent | 165 (73.3) | 142 (66.7) | 0.128 |
| Present | 60 (26.7) | 71 (33.3) | |
| **Tumour budding** | | | |
| Low | 168 (74.3) | 134 (64.1) | 0.021 |
| High | 58 (25.7) | 75 (35.9) | |
| **Mucin Lakes** | | | |
| Absent | 95 (80.5) | 41 (59.4) | 0.002 |
| Present | 23 (19.5) | 28 (40.6) | |
| **Differentiation** | | | 0.722 |
| Well differentiated | 202 (88.6) | 189 (87.5) | |
| Poor/moderate differentiation | 26 (11.4) | 27 (12.5) | |
| **Peritoneal Involvement** | | | |
| Absent | 185 (80.4) | 156 (71.9) | 0.034 |
| Present | 45 (19.6) | 61 (28.1) | |
| **mGPS** | | | |
| 0 | 149 (64.8) | 126 (58.1) | 0.112 |
| 1 | 44 (19.1) | 45 (20.7) | |
| 2 | 37 (16.1) | 46 (21.2) | |
| **GMS** | | | |
| 0 | 40 (17.8) | 30 (14.4) | 0.441 |
| 1 | 142 (63.1) | 136 (65.4) | |
| 2 | 43 (19.1) | 42 (20.2) | |

**Table S1. Clinical features of CRC TMA under comparison for PODXL Low versus High expression.**
